# SOX transcription factors direct TCF-independent WNT/beta-catenin transcription

**DOI:** 10.1101/2021.08.25.457694

**Authors:** Shreyasi Mukherjee, David M. Luedeke, Leslie Brown, Aaron M. Zorn

## Abstract

WNT/ß-catenin signaling regulates gene expression across numerous biological contexts including development, stem cell homeostasis and tissue regeneration, and dysregulation of this pathway has been implicated in many diseases including cancer. One fundamental question is how distinct WNT target genes are activated in a context-specific manner, given the dogma that most, if not all, WNT/ß-catenin responsive transcription is mediated by TCF/LEF transcription factors (TFs) that have similar DNA-binding specificities. Here we show that the SOX family of TFs direct lineage-specific WNT/ß-catenin responsive transcription during the differentiation of human pluripotent stem cells (hPSCs) into definitive endoderm (DE) and neuromesodermal progenitors (NMPs). Using time-resolved multi-omics analyses, we show that ß-catenin association with chromatin is highly dynamic, colocalizing with distinct TCFs and/or SOX TFs at distinct stages of differentiation, indicating both cooperative and competitive modes of genomic interactions. We demonstrate that SOX17 and SOX2 are required to recruit ß-catenin to hundreds of lineage-specific WNT-responsive enhancers, many of which are not occupied by TCFs. At a subset of these TCF-independent enhancers, SOX TFs are required to both establish a permissive chromatin landscape and recruit a WNT-enhanceosome complex that includes ß-catenin, BCL9, PYGO and transcriptional coactivators to direct SOX/ß-catenin-dependent transcription. Given that SOX TFs are expressed in almost every cell type, these results have broad mechanistic implications for the specificity of WNT responses across many developmental and disease contexts.

## INTRODUCTION

WNT/ß-catenin signaling is used reiteratively in all metazoans with critical roles throughout the life of an organism ranging from cell-fate determination in embryogenesis, organogenesis and tissue regeneration to adult stem-cell homeostasis^1^. Dysregulation of the WNT pathway is associated with a range of human diseases, from cancer to neurodegeneration^2^. Despite being the subject of intense study for decades, how the WNT pathway executes its context-dependent roles through the selective transcription of distinct context-specific target genes remains poorly understood ^3, 4^.

In the canonical WNT pathway, recruitment of ß-catenin (CTNNB1) to enhancers is the key event initiating transcription^5–7^. In cells not receiving a WNT signal, cytosolic ß-catenin is phosphorylated by glycogen synthase kinase (GSK3ß) and targeted to a proteosomal degradation complex. The binding of WNT ligands to FZD/LRP receptors on the cell surface leads to the inactivation of the destruction complex, allowing non-phosphorylated ß-catenin to accumulate and localize to the nucleus. Upon its translocation to the nucleus, ß-catenin associates with DNA-binding TCF/LEF (hereafter TCF) transcription factors (TFs), where it is thought to displace, and/or lead to the inactivation of, TLE/Groucho co-repressors resulting in the transcription of WNT target genes^3, 8^. Recent studies have provided a more integrated view of this transcription complex, known as the “WNT-enhanceosome”, that is assembled on WNT responsive enhancers (WREs). ß-catenin/TCF on chromatin interact with several cofactors including: BCL9, PYGOPUS, CHIP/LDB/SSDP and the BAF complex. Current models propose that upon WNT signaling, recruitment of ß-catenin results in a conformational change in the WNT-enhanceosome and an association of WREs with their cognate promoters, recruitment of histone acetyltransferases (Ep300/CBP), activation of RNA polymerase II and transcription initiation^9–11^. Yet, how ß-catenin is recruited to distinct WREs in different cellular contexts remains a mystery. One of the major challenges is that ß-catenin itself cannot bind DNA and is dependent on interactions with DNA-binding TFs for its recruitment to WREs^5^.

Decades of research indicate that the TCF family of HMG-box TFs are core mediators of WNT-responsive transcription by physically interacting with ß-catenin and recruiting it to WREs. There are four mammalian TCFs: TCF7, TCF7L1, TCF7L2 and LEF1^12^ that *in vitro* all bind as monomers to nearly identical 5’-SATCAAGS-3’ DNA-sequences^13^. Genetic studies indicate that different TCFs have largely redundant functions^14, 15^ and ChIP-Seq datasets show that they co-occupy many of the same chromatin loci *in vivo*^16, 17^. A few mechanisms have been proposed to account for some degree of variability in DNA-binding, such as alternative RNA-splicing variants of TCF^18^, but this is insufficient to account for the diversity of WNT target genes. Thus, it remains largely unclear how TCFs alone with very similar DNA-binding specificities can regulate the vast diversity of context specific WNT-responsive transcription. Over the years, increasing evidence has emerged that ß-catenin can interact with other TFs in addition to TCFs, including OCT4, TBX3, HIF1*α*, SMAD and SOX TFs^19–22^, however whether these can recruit ß-catenin to context-specific enhancers throughout the genome to account for the diversity of WNT-response transcription is unclear.

One of the strongest candidates for an alternative TF family that could mediate Wnt-responsive transcription is the SOX family. Related to TCFs, the twenty SOX factors in humans are conserved HMG-box DNA-binding domain TFs that recognize similar but distinct variations of a core WWCAAW motif^23–25^. Many SOX TFs have been reported to bind to ß-catenin *in vitro* and can antagonize or potentiate WNT-responsive transcription of an artificial TOP:flash reporter, in overexpression conditions^26–34^. However, relatively little is known about SOX/ß-catenin interactions *in vivo* and the extent to which SOX TFs can modulate the specificity of WNT responsive transcription. In mouse neural progenitor cells, Sox2 can interact with ß-cat/Lef1 to repress the well-known pro-proliferative Wnt target gene *Cyclin D1*^35^. Moreover, we recently showed in *Xenopus* gastrulae that Sox17 and ß-catenin co-occupy endodermal enhancers to regulate lineage-specific gene expression^27, 36^. However, whether Sox17 mechanistically regulates ß-catenin chromatin association remains unclear.

In this study, we test the hypothesis that SOX TFs can function as lineage-specific mediators of WNT/ß-catenin transcription during development. Using the directed differentiation of human pluripotent stem cells (hPSCs) into definitive endoderm (DE) and neuromesodermal progenitors (NMPs) and time-resolved ChIP-Seq, ATAC-Seq and RNA-Seq experiments, we show that ß-catenin chromatin binding is highly dynamic and is associated with distinct genomic loci as differentiation proceeds. We show that ß-catenin colocalizes with different combinations of just TCFs, TCF and SOX, or just SOX TFs, with evidence of both cooperative and competitive genomic interactions. We demonstrate that SOX17 (in DE) and SOX2 (in NMPs) are required to recruit ß-catenin to hundreds of lineage-specific WNT-responsive enhancers, many of which are not bound by TCFs. At a subset of these TCF-independent enhancers, SOX TFs are required to both establish a permissive chromatin landscape and recruit a WNT-enhanceosome complex including ß-catenin, BCL9, PYGO and transcriptional coactivators to direct SOX/ß-catenin-dependent transcription. These results have important implications for how the combination of TCFs and lineage-specific TFs such as the SOXs might regulate distinct transcriptional programs downstream of WNT signaling in diverse biological contexts.

## RESULTS

### ß-catenin binds dynamically to distinct lineage-specific enhancers during endoderm differentiation

To characterize lineage-specific genomic recruitment of ß-catenin during development, we performed a time course analysis of hPSCs differentiated to DE. We treated hPSCs with recombinant Activin A and the GSK3ß inhibitor CHIR99021 to stimulate the WNT/ß-catenin pathway for three days [Fig 1A]. This protocol mimics the NODAL and WNT signaling in the gastrula embryos and generated primitive streak-like mesendoderm cells on Day 1, endoderm progenitors on Day 2 and a highly homogenous [>90%] population of SOX17-expressing DE cells on Day 3 [Fig 1A, B]. Immunostaining and western blot analyses confirmed that total ß-catenin levels and the transcriptionally active, K49-acetylated isoform of ß-catenin^37^ was present in the nucleus of all cells from Days 1-3 [Fig 1B and S1A, B]. We then performed chromatin immunoprecipitation coupled with high-throughput sequencing (ChIP-Seq) each day to profile ß-catenin chromatin binding during the progressive differentiation from pluripotency to DE. In the Day 0 ‘WNT-OFF’ pluripotency state, we detected negligible ß-catenin binding with <750 peaks, which increased upon CHIR treatment to 4501 peaks at Day1, 18608 peaks on Day2 and 11864 peaks on Day3 [Fig 1C, D]. Interestingly, ß-catenin binding was observed at distinct genomic regions on different days, suggesting that it regulated distinct transcriptional programs as lineage specification proceeded. To identify these dynamic chromatin binding patterns, we merged all significantly called peaks from all days and performed k-means clustering [Fig 1C, F-J; see Methods]. This identified five distinct patterns: common peaks shared across all time points [n = 1972], Day 1 enriched peaks [n = 1304], Day 2 enriched peaks [n = 8364], peaks specific to Days 2 and 3 [n = 7065]; and Day 3 enriched peaks [n = 2611] [Fig 1C, F-J].

**Figure 1.**
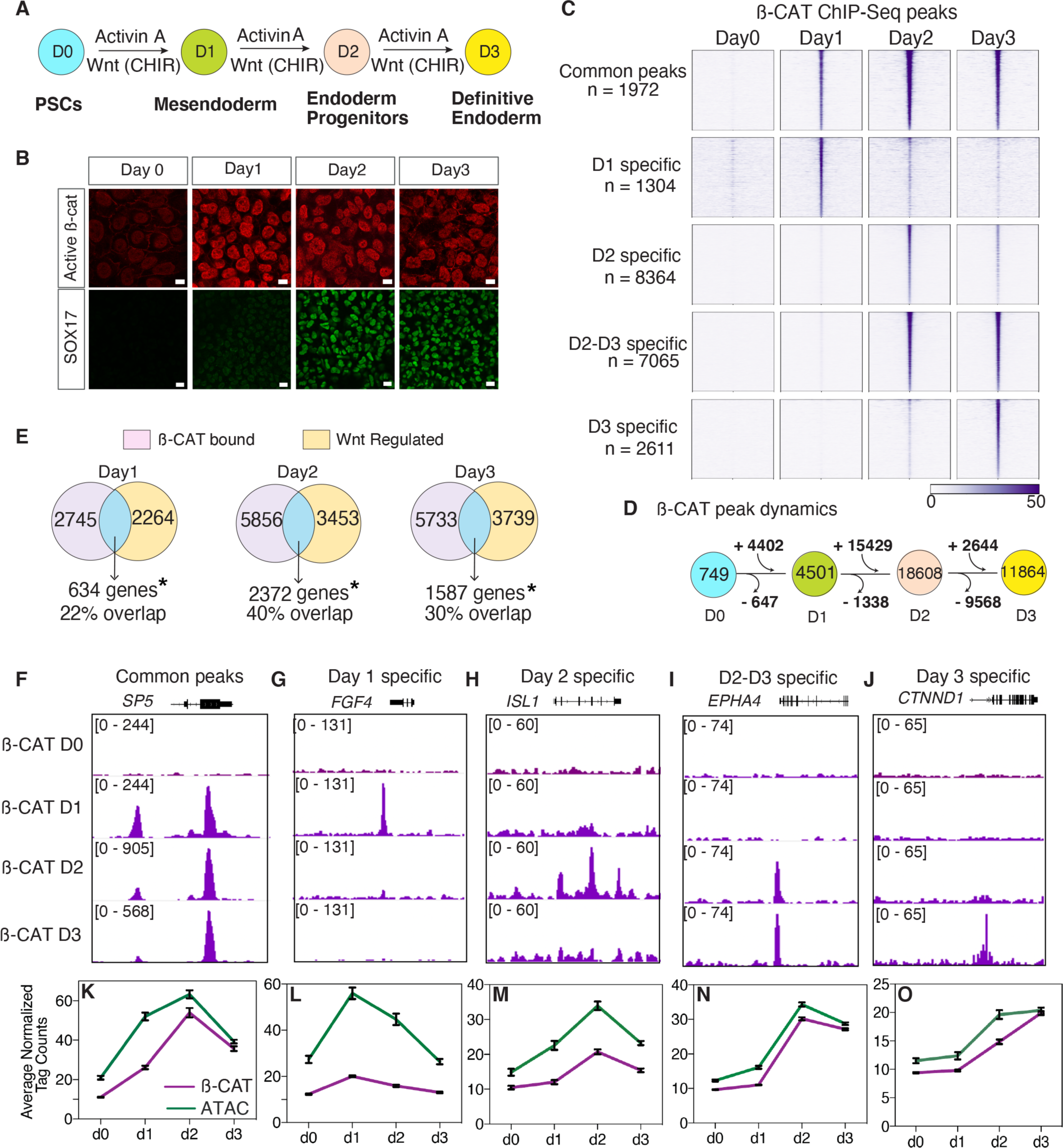
Dynamic chromatin binding of ß-catenin during DE differentiation. **A.** Schematic of definitive endoderm differentiation. **B.** Immunostaining of nuclear K49ac ‘active’ and SOX17 at days 0 - 3 (scale bar = 20 and 50 μm respectively) **C.** Density plot of ß-catenin ChIP-seq showing five categories of temporally distinct peaks. **E.** Overlap of Wnt regulated genes with genes associated with ß-catenin bound peaks. *Significant overlap based on hypergeometric test, day 1: *p* = 4.75 × 10^-105^; day 2: *p =* 3.63 × 10^-138^; day 3: *p* = 2.52 × 10^-48^. **D.** Schematic of ß-catenin peak dynamics during differentiation. **F – J.** Genome browser tracks showing ß-catenin chromatin binding for each of the five categories of peaks. **K – O.** Average normalized ß-catenin (purple) and ATAC-Seq (green) read density, plotted as a line graph. Error bars represent standard error of mean for each category.

We next sought to identify which ß-catenin binding events were associated with WNT-responsive transcription. We performed RNA-Seq experiments every 24 hours during the differentiation trajectory with or without WNT stimulation; cells were treated with Activin for 3 days with either a WNT agonist (CHIR99021) or a WNT antagonist (C59) [Fig S2A, see Methods]. Differential expression analysis of the WNT-ON and WNT-OFF states identified WNT-responsive transcripts on each day [Fig S2B – E, see Methods]. Principal component analysis showed that when WNT signaling was inhibited from the beginning, cells could not differentiate and retained a pluripotency-like transcriptional signature [Fig S2B] demonstrating that WNT is required for DE differentiation. Integrating the ChIP-Seq and RNA-Seq data confirmed that the dynamic genomic binding of ß-catenin was indeed associated with different transcriptional programs on different days [Fig 1D]. Unsupervised hierarchical clustering of all the direct WNT activated genes associated with ß-catenin binding across all days [Fig S2F–I] identified several distinct groups of coregulated genes similar to the dynamic ß-catenin binding patterns. These included ‘common’ WNT-responsive genes activated by CHIR irrespective of the day of differentiation [n=130] as well as genes specifically regulated by CHIR on Day 1 [n = 121], Day 2 [n = 857] and Day 3 [n = 338] [Fig S2F – I]. Gene ontology (GO) enrichment analysis [Fig S1C-G, S2F-I] revealed that “common” genes associated with ß-catenin binding and CHIR-dependent expression at all time points included well-known universal WNT targets such as *SP5* and *AXIN2* and were associated with terms such as ‘Wnt pathway’ and ‘cell-fate specification’ [Fig S1C-G, S2F-1]. In contrast, direct ß-catenin targets on Days 1-2 were enriched for terms related to gastrulation, mesoderm, and endoderm formation. These included primitive streak genes such as *FGF4*, *ISL1* and *EPHA4* which are expressed in both the mesoderm and endoderm. Day 3 direct targets were frequently associated with genes expressed in mature epithelium such as *CTNND1* [Fig 1F-J, S2I]. Therefore, progressive DE specification is associated with rapid and dynamic ß-catenin relocalization to distinct lineage-specific genomic loci.

### Changes in chromatin accessibility is not sufficient to account for dynamic ß-catenin binding

One possible explanation for the dynamic binding of ß-catenin to distinct genomic loci was changes in the underlying chromatin accessibility landscape.^38^ For example, Day 1 specific ß-catenin bound loci may only be accessible on Day 1 but compacted into nucleosomes on Days 2-3. To test this hypothesis, we performed time-course ATAC-Seq experiments every 24 hours during differentiation and compared this with ß-catenin binding [Fig 1K-O; Fig S1H - V]. This revealed that although newly gained ß-catenin binding sites were associated with increased chromatin accessibility at lineage-specific loci [Fig 1K-O, Fig S1M-Q], in general, most ß-catenin-bound loci were accessible at all days of differentiation [Fig S1 R-V]. This was particularly exemplified in peaks enriched specifically at Days 1 and 2, where ß-catenin binding was rapidly lost as differentiation proceeded despite the fact that most of the chromatin remained accessible; >60% on Day 2 and > 40% on Day 3 [Fig S1 N,S]. Altogether, this time-resolved genomic analyses of ß-catenin occupancy indicates that lineage-specific WNT responsive transcription is mediated by rapid ß-catenin relocalization, highlighting further the need to understand the role of TFs in mediating this recruitment.

### SOX17 and ß-catenin co-occupy an increasing number of genomic sites during endoderm differentiation

Next, we sought to understand the extent to which dynamic localization of ß-catenin to different genomic regions over time could be explained by TCFs or SOX17, the main SOX TF regulating endoderm development. Examination of the RNA-seq data showed that all four TCF/LEFs were expressed during DE differentiation [Fig 2A]. Consistent with previous studies^39–,41^, *TCF7L1* and *TCF7L2* were the most highly expressed TCFs in pluripotent cells, whereas *TCF7* and *LEF1* were not expressed in pluripotent cells but activated in response to WNT during differentiation. We performed ChIP-Seq experiments every 24 hours to profile the binding dynamics of all four TCFs and SOX17 during DE differentiation and compared this to ß-catenin occupancy. Peak overlap analysis revealed that 96% of ß-catenin binding events [n = 4328/4501] in Day 1 cells could be accounted for by occupancy of at least one TCF, consistent with the concept of TCFs as predominant mediators of ß-catenin binding [Fig 2B].

**Figure 2.**
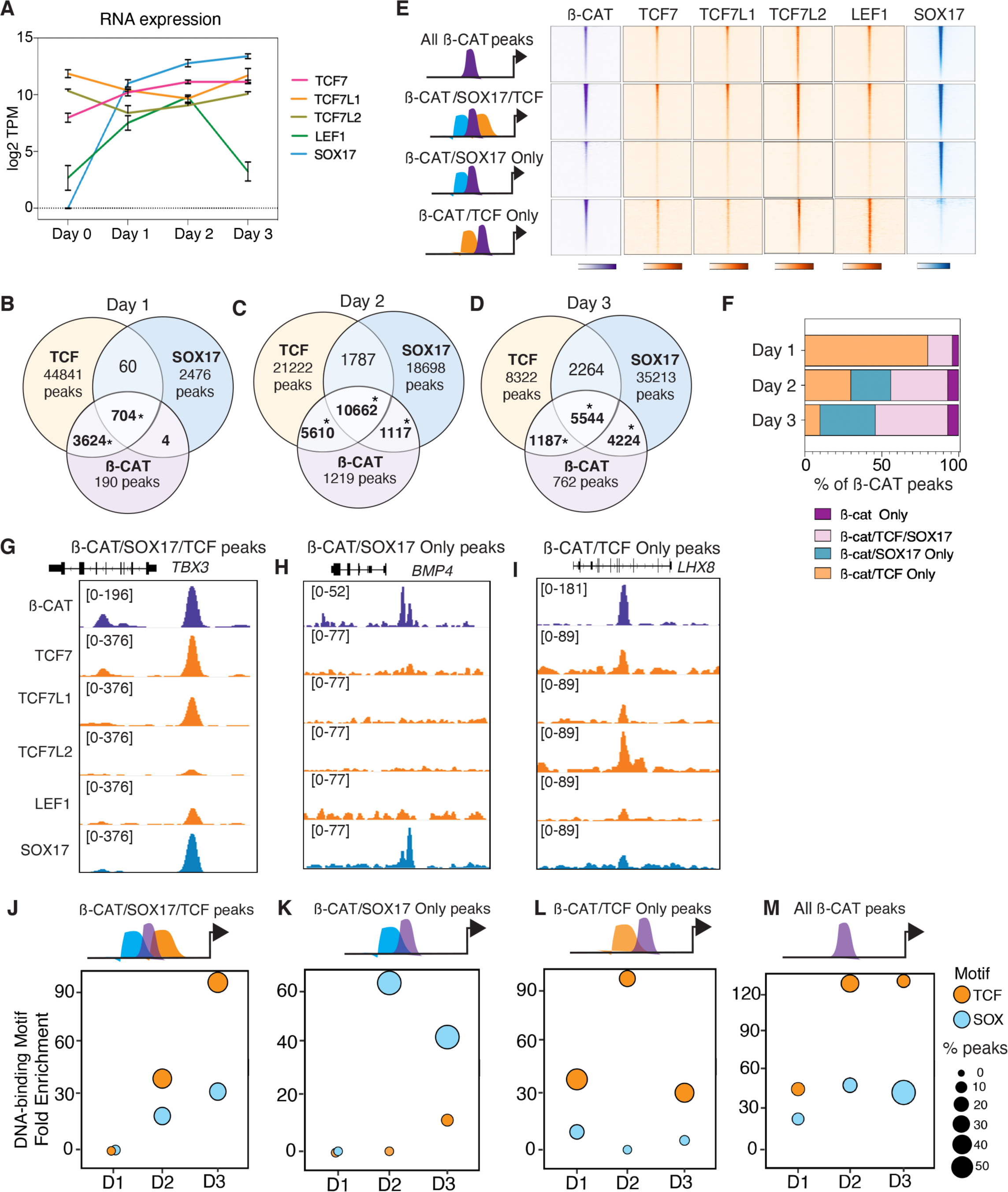
Dynamic co-localization of ß-catenin with SOX17 and TCFs. **A.** RNA-seq expression levels of *SOX17, TCF7, TCF7L1*, *TCF7L2* and *LEF1* during endoderm differentiation. **B – D.** Venn diagrams showing peak overlap of ß-catenin, TCFs, and SOX17 during each day of endoderm differentiation. *Significant overlap based on hypergeometric test, p<0.0001. **E.** ChIP-seq density plots showing ß-catenin, TCF and SOX17 co-occupancy on Day 3 at four categories of loci: All Day 3 ß-catenin peaks, ß-catenin/SOX17/TCF cobound peaks, ×-catenin/SOX17 only peaks, ß-catenin/TCF only peaks. **F.** Stacked bar graph plotting the percentage peak overlap of ß-catenin with TCFs and/or SOX17. **G – I.** Representative genome browser views of genes with associated co-binding of ß-catenin with SOX17 and TCF **(G)**, ß-catenin and SOX17 only **(H)** and ß-catenin and TCF only **(I)** peaks. **J – M.** Scatter plots showing fold enrichment and proportion of peaks containing SOX or TCF DNA-binding motifs in different categories of ß-catenin peaks across differentiation.

Surprisingly however, as differentiation progressed, we observed a progressive shift in co-localization of ß-catenin from TCFs to SOX17. In Day 2 cells, TCFs colocalized with ß-catenin at 87% [n = 16272/18608] of genomic loci, and by Day 3, TCFs accounted for only 57% of ß-catenin binding [n = 6731/11864] [Fig 2B-D, F, Fig S3A-F]. In contrast, the number of loci co-occupied by ß-catenin and SOX17 increased from 16% at Day 1 to 43% at Day 2 and 83% in Day3 DE cells [Fig 2B-D, F, Fig S3A-F]. Focusing on Day 3 DE, we identified four categories of peaks [Fig 2E, F]: those co-occupied only by ß-catenin and SOX17 but not TCFs such as that associated with *BMP4* [n = 4224/11717, 36%], loci bound by ß-catenin/SOX17/TCF such as *TBX3* [n = 5544/11717, 47%], loci co-bound by ß-catenin and at least one TCF, but not bound by SOX17 including *LHX8* [n = 1187/11717, 10%] and ß-catenin binding events where we could not detect co-binding of either TCFs or SOX17 [n = 762/11717, 7%] [Fig 2E-I]. *De-novo* motif analysis revealed, as expected for DE enhancers^42^, an enrichment of GATA, FOXA and SOX DNA-binding motifs across all Day 3 peak categories [Fig S3G]. Time-resolved analyses of TCF and SOX motif enrichment in the different categories showed an increase of SOX motifs relative to TCF DNA binding sites that correlated with increased SOX17/ß-catenin co-occupancy [Fig 2J – M]. These data argue that TCFs alone cannot account for the total extent of ß-catenin recruitment to chromatin during progressive differentiation and indicate that DE-specific ß-catenin binding events are correlated with co-occupancy of SOX17 or SOX17 and TCFs.

### SOX17 is required to recruit ß-catenin to DE specific WNT-responsive enhancers

We next assessed whether SOX17 was required to recruit ß-catenin to chromatin, particularly in genomic regions not co-occupied by TCFs. To test this, we generated a CRISPR-mediated SOX17-null mutant iPSC line [Fig S4A] and asked whether recruitment of ß-catenin to DE-specific genomic loci was compromised. Immunostaining and western blots confirmed a loss of SOX17 and showed that total and nuclear ß-catenin protein levels were unaltered in SOX17 knockout (KO) cells [Fig S4 B, C]. Similarly, loss of SOX17 did not affect mRNA or protein levels of TCF7, TCF7L1 or TCF7L2, however, LEF1 was upregulated [Fig S4 D-F]. Immunostaining for FOXA2 as well as RNA-Seq analysis confirmed that DE specification was compromised in SOX17KO cells [Fig S4B, G - K].

We next performed ß-catenin ChIP-Seq on Day 3 wildtype [WT] and SOX17KO cells, identifying 13,131 peaks bound by ß-catenin in WT cells as opposed to 41,058 ß-catenin peaks in SOX17KO cells. Differential binding analysis revealed three distinct categories of ß-catenin binding: i) ß-catenin peaks lost in SOX17KO such as that associated with *BMP7* [n = 4337], ii) ß-catenin peaks largely unchanged in WT and SOX17KO like *DKK1* [n = 3894] and iii) ß-catenin peaks gained in SOX17KO cells such as *MEOX1* [n = 20496] [Fig 3A-F, Fig S5 A-D]. We next assessed the extent to which SOX17-dependent changes in ß-catenin binding correlated with TCF co-occupancy [Fig 3A-C] by performing ChIP-Seq for all TCFs in both WT and SOX17KO cells. We then quantified the status of SOX17 and TCF co-occupancy for each of these three groups of ß-catenin peaks by differential occupancy analysis. This revealed that 96% [n = 4136/4337] of ß-catenin binding events lost in SOX17KO cells were co-bound by SOX17 in WT cells, with only 30% [n = 1318/4337] also being co-bound by TCFs. Thus, the majority of the ß-catenin peaks lost in the SOX17KO cells showed no evidence of TCF co-occupancy. In contrast, 65% [n = 2088/3894] of the ß-catenin peaks that were unchanged between WT and SOX17KO were co-occupied by TCFs in WT cells and this increased to 90% TCF co-occupancy in SOX17KO cells. The striking increase in *de-novo* ß-catenin binding events in SOX17KO cells was also associated with an increase in TCF-co-occupancy from 18% [n = 4373/24096] in WT DE to 85% [n = 20008/24096] in SOX17KO cells as exemplified by the mesoderm-specific gene *MEOX1* [Fig 3C, F]. Interestingly, GO enrichment analysis indicated that genes associated with lost ß-catenin peaks were enriched for endoderm organogenesis, whereas genes associated with gained ß-catenin peaks were enriched for cardiac mesoderm and epithelial to mesenchymal transition [Fig. S5I]. These data indicate that in the absence of SOX17, TCFs (primarily TCF7L2 and LEF1), recruit ß-catenin to different enhancers, many of which are associated with alternative lineages [Fig S5 G,H].

**Figure 3.**
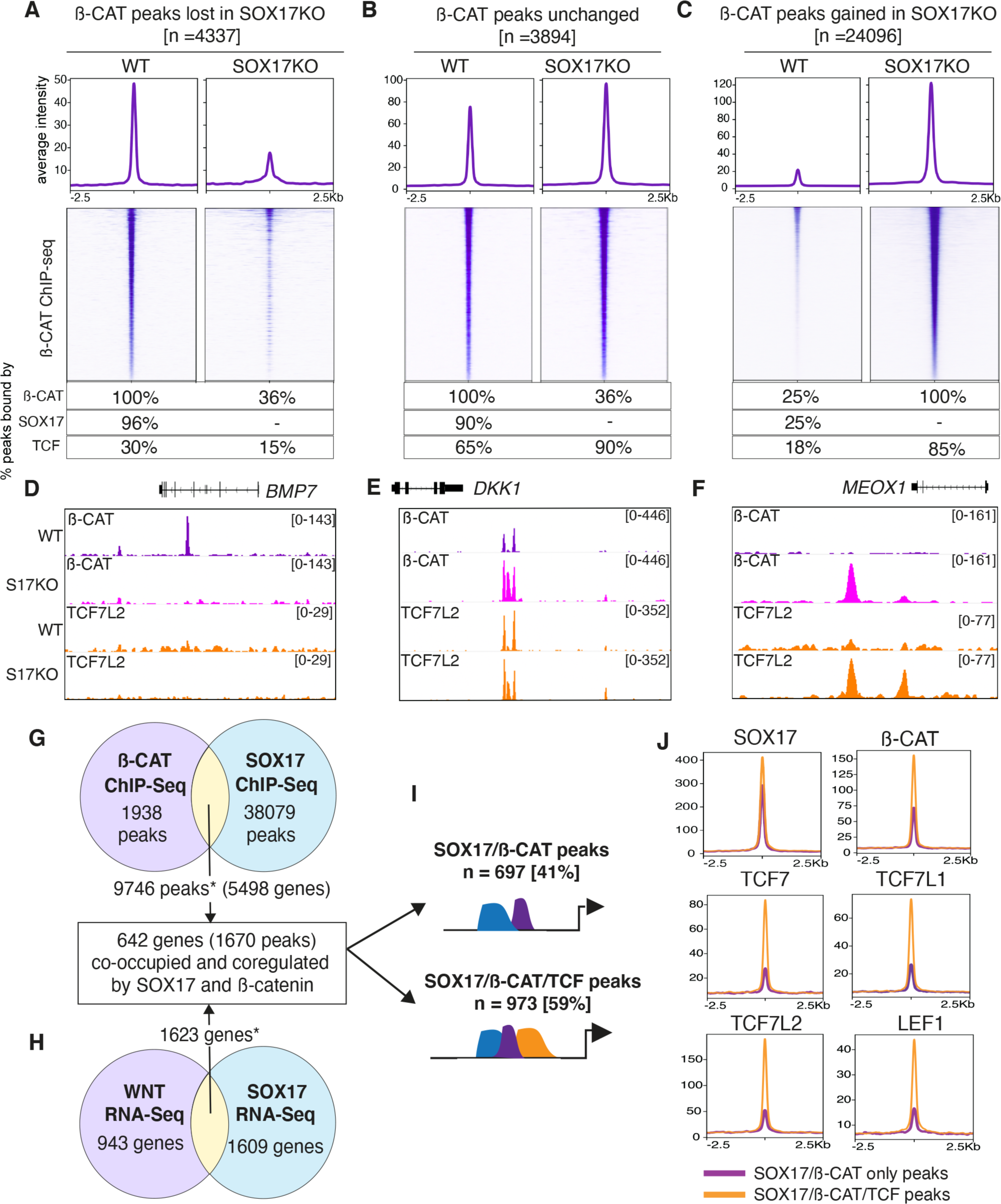
SOX17 is required for ß-catenin recruitment to chromatin in the absence of TCFs. **A-C** ß-catenin ChP-seq density plots and metaplots of the average signal intensity for three distinct categories of peaks: **A.** ß-catenin peaks lost in SOX17KO cell, **B**. ß-catenin peaks that remain unchanged between WT and SOX17KO and cells and **C.** new ß-catenin peaks gained in SOX17KO cells. The tables below each density plot show the percentage of peaks bound by ß-catenin, SOX17 or any TCF. **D – F.** Genome browser view showing ß-catenin and TCF7L2 chromatin occupancy in WT and SOX17KO cells at representative loci for each category of peaks. **G - H.** Integration of ß-catenin and SOX17 RNA-Seq and ChIP-Seq datasets from Day 3 defining direct co-regulated genes. * Significant overlap based on hypergeometric test, ChIP RNA-Seq peak overlap: *p* = 1.03 × 10^-553^; -Seq gene set overlap: *p* = 2.07 × 10^-61^. **I.** Diagram showing percentage of SOX17/ß-catenin coregulated that peaks are also co-bound by TCFs or not. **J.** ChIP-seq metaplots showing the average peak intensity for SOX17, ß-catenin, TCF7, TCF7L1, TCF7L2 and LEF1 at both categories of SOX17/ß-catenin coregulated peaks: SOX17/ß-catenin only (purple) and SOX17/ß-catenin/TCF cobound peaks (orange).

We next determined the extent to which these changes in ß-catenin binding in SOX17KO cells was associated with changes in ß-catenin/SOX17-dependent transcription. We performed RNA-Seq on Day 3 WT and SOX17KO cells [FigS4G], identifying 3232 differentially regulated transcripts. GO analysis of transcripts upregulated in SOX17KO [n = 1424] revealed enriched terms related to mesoderm whereas downregulated transcripts were enriched for endoderm differentiation terms [Fig S4H,I, Fig S6C]. Integrating the SOX17 and ß-catenin ChIP-Seq and RNA-Seq datasets, we identified 642 genes [corresponding to 1670 peaks] that were coordinately co-occupied and coregulated by SOX17 and ß-catenin [Fig 3G, H, Fig S6A, B]. Epigenetic analysis showed that these SOX17/ß-catenin cobound peaks were enriched for H3K4me1 and H3K27ac, histone marks indicative of poised and active enhancers [Fig S6D]^43, 44^. Further analysis revealed that SOX17/ß-catenin co-activated genes had almost exclusively endoderm enriched expression, whereas a substantial number of the SOX17/ß-catenin repressed and SOX17 repressed/ß-catenin activated genes were enriched in mesectodermal lineages [Fig S6C]. This was consistent with our recent finding in *Xenopus* gastrulae where Sox17 promotes endoderm fate while repressing mesectoderm identity^36^. Next, we investigated the extent to which these SOX17/ß-catenin coregulated and co-occupied enhancers were bound by TCFs. This revealed that 41% of SOX17/ß-catenin coregulated enhancers had little to no evidence of TCF binding [Fig 3I, J, Fig S6E].

Together these data demonstrate that SOX17 is required to recruit ß-catenin to a subset endoderm-specific WNT-responsive enhancers, many of which have no evidence of TCF binding. Loss of SOX17 leads to widespread relocalization of ß-catenin genomic binding, in many cases recruitment to mesodermal enhancers by TCFs, suggesting that SOX17 and TCFs might compete to recruit ß-catenin to different lineage-specific loci [Fig S6F].

### SOX17 is required to establish a permissive chromatin landscape at a subset of TCF-independent endodermal enhancers

There is evidence that SOX TFs can act as pioneering factors by directly engaging nucleosomes to regulate chromatin accessibility^45–48^, which might explain in part the loss of ß-catenin binding in SOX17KO cells. To address this possibility, we performed ATAC-Seq experiments in Day 3 WT and SOX17KO cells. Differential peak analysis revealed 13737 genomic regions with significantly increased accessibility based on ATAC-seq and 29580 regions that were significantly less accessible in SOX17KO cells [Fig S7A]. We focused our analysis on those enhancers that lost ß-catenin binding in SOX17KO cells and that were enriched for SOX17 but not TCF occupancy; we termed these ‘TCF-independent’ enhancers [Fig 4]. About half of the TCF-independent enhancers [52%, n = 1577/3020] displayed decreased chromatin accessibility in SOX17KO cells such as *SALL1* [termed as Class I enhancers] while the others did not display significant SOX17-dependent changes in accessibility like *PRIMA1* [termed as Class II enhancers] [Fig 4A, F]. We used the Nucleoatac package^49^ to predict nucleosome occupancy at both classes of enhancers. We observed a similar dip in nucleosome occupancy at the ATAC-seq peak centers of both Class I and Class II enhancers in WT cells, but in SOX17KO cells, there was a significant increase in nucleosome occupancy in Class I relative to Class II enhancers, confirming that SOX17 is required to establish chromatin accessibility at about half of the TCF-independent enhancers [Fig 4B, C].

**Figure 4.**
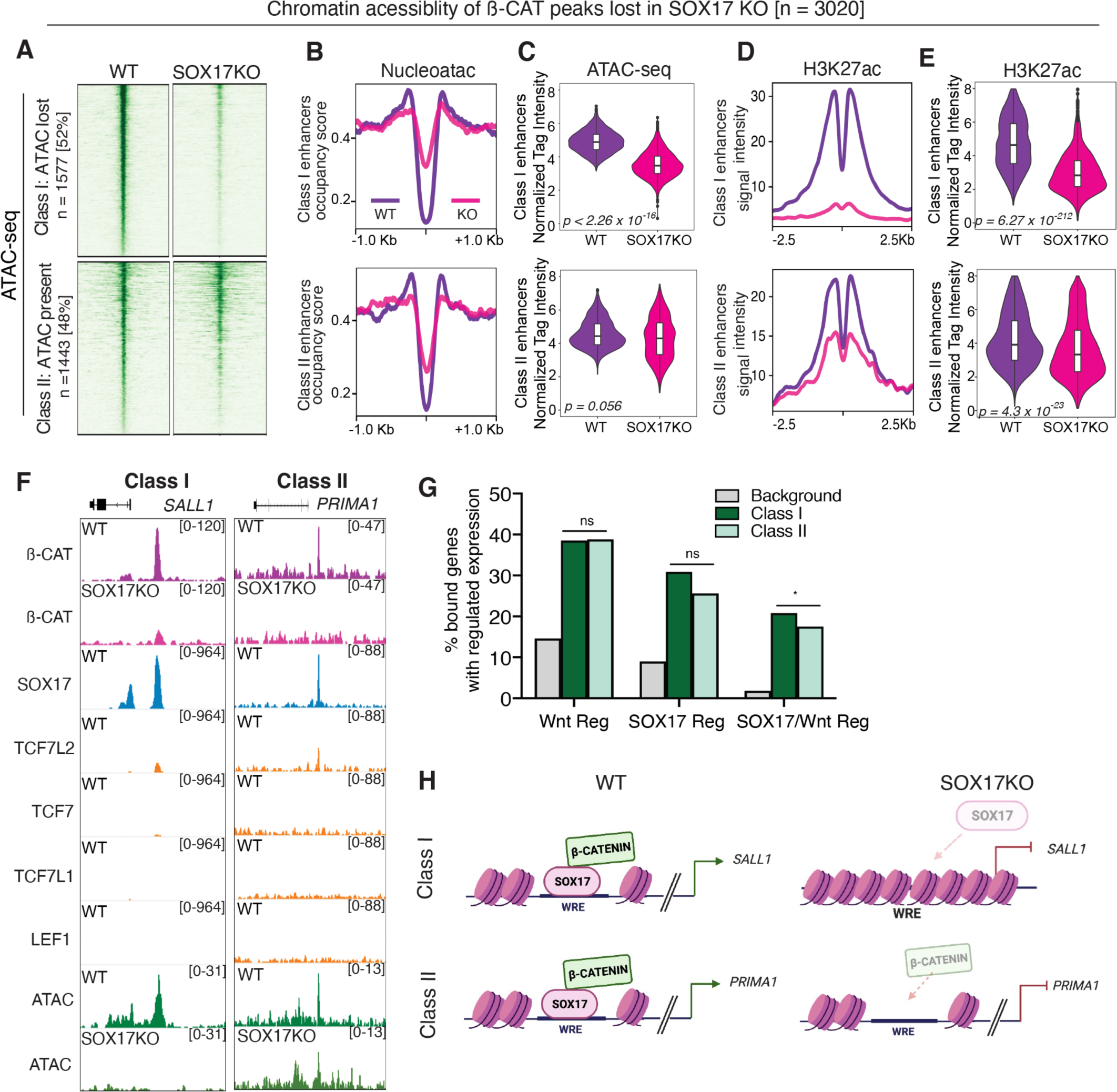
Chromatin accessibility only partially accounts for loss of ß-catenin binding in SOX17 KO cells. **A.** Density plots showing ATAC-Seq signal intensity in WT and SOX17KO cells for two classes of SOX17-dependent ß-catenin bound peaks. Class I peaks with reduced accessibility and Class II enhancers which loose ß-catenin in SOX17 KO cells but accessibility is unchanged. **B.** Metaplots showing nucleosome occupancy signals and **C**. quantification of ATAC-Seq read densities in WT (purple) and SOX17KO (pink) cells for both Class I and Class II enhancers**. D.** Metaplots showing the average H3K27ac ChIP-seq signal for Class I and Class II enhancers and E. violin plots quantifying tag density. Indicated p-values were determined by Wilcoxon rank sum test. **F.** Genome browser view of representative Class I and Class II enhancers. **G.** Bar graph showing the proportion of Class I and Class II enhancers associated with Wnt regulated, SOX17 regulated and SOX17/Wnt coregulated genes. Fisher’s exact test, comparing proportion of Class I vs. Class II enhancers associated with regulation. *p* = 0.88 for Wnt regulated genes, *p* = 0.472 for SOX17 regulated genes, *p* = 0.02 for SOX17/Wnt coregulated genes. Background is all genes in the genome. **H.** Schematic of SOX17 and ß-catenin recruitment to Class I and Class II Wnt responsive enhancers (WRE).

We next investigated whether SOX17 was required for the deposition of the histone mark H3K27ac, a signature of transcriptionally active enhancers^50^. H3K27ac ChIP-Seq of Day 3 WT and SOX17KO cells showed a significant loss of H3K27ac deposition in SOX17KO cells at both Class I and Class II enhancers, albeit to a lesser extent on Class II [Fig 4D, E]. Consistent with this, analysis of gene expression associated with Class I and Class II enhancers revealed that they were coregulated by both SOX17 and ß-catenin to a similar extent [Fig 4G, Fig S7B], suggesting that loss of chromatin accessibility alone is not sufficient to account for the failure to activate these Wnt-responsive enhancers.

A similar analysis of peaks with unchanged ß-catenin binding in WT and SOX17KO showed little, if any, changes in chromatin accessibility or SOX17-dependent H3K27ac deposition [Fig S7C – G]. In contrast, of the loci that gained *de-novo* ß-catenin and TCF binding in SOX17KO cells, 59% [n = 14291/24096] exhibited an open chromatin signature in SOX17KO cells. Of these, 67% [n = 7908/14291] exhibited significantly increased chromatin accessibility in SOX17KO whereas the rest were unchanged [Fig S7H - J]. In both cases there was elevated H3K27ac deposition at these loci in the absence of SOX17, [Fig S7K, L] consistent with activation of a mesoderm transcriptional program.

Collectively, these results indicate that SOX17 can regulate lineage specific Wnt-responsive transcription both by regulating chromatin accessibility and by recruiting ß-catenin to a subset of TCF-independent endoderm-specific enhancers. In addition, the data suggests that upon loss of SOX17, mesoderm-specific loci become accessible [Fig S4J, K, Fig S7H-L] and that TCFs can then recruit ß-catenin to activate alternative Wnt-responsive transcriptional programs at these WREs.

### Lineage-specific recruitment of ß-catenin is a general feature of SOX TFs

Next, we investigated whether other SOX TFs also had the ability to regulate ß-catenin chromatin binding and lineage-specific WNT-responsive transcription. To test this, we used hPSC-derived bipotent neuromesodermal progenitors (NMPs) where antagonistic interactions between SOX2 and the TF TBXT control WNT-responsive cell fate decision between neural and mesoderm-derived lineages^51–53^. Previous ChIP-Seq analysis of mouse NMPs differentiated from ESCs has shown that SOX2 co-occupies enhancers of several NMP targets together with TBXT and ß-catenin^53^.

To investigate SOX2/ß-catenin interactions in NMPs, we used a CRISPRi SOX2 iPSC line where deactivated Cas9 fused to the KRAB repressor domain is used to repress *SOX2* expression in a doxycycline-dependent manner^54^. Using previously published protocols^55^, we directed the differentiation of PSCs towards the NMP lineage through the addition of FGF8b and the WNT agonist CHIR99021^55^, generating relatively pure (>70%) populations of NMPs co-expressing SOX2 and TBXT after 3 days of culture [Fig 5A,B]. To knock down (KD) SOX2 levels, we treated cells with dox right after exit from pluripotency but before the onset of NMP specification [Fig 5A, see Methods]. Immunostaining confirmed loss of SOX2 in the KD cells and showed that there was no change in overall levels total or active nuclear ß-catenin [Fig 5CB, Fig S8BG]. Moreover, we did not observe any significant difference in mRNA or protein levels of any of the TCFs in WT versus SOX2KD cells [Fig S8G, H].

**Figure 5.**
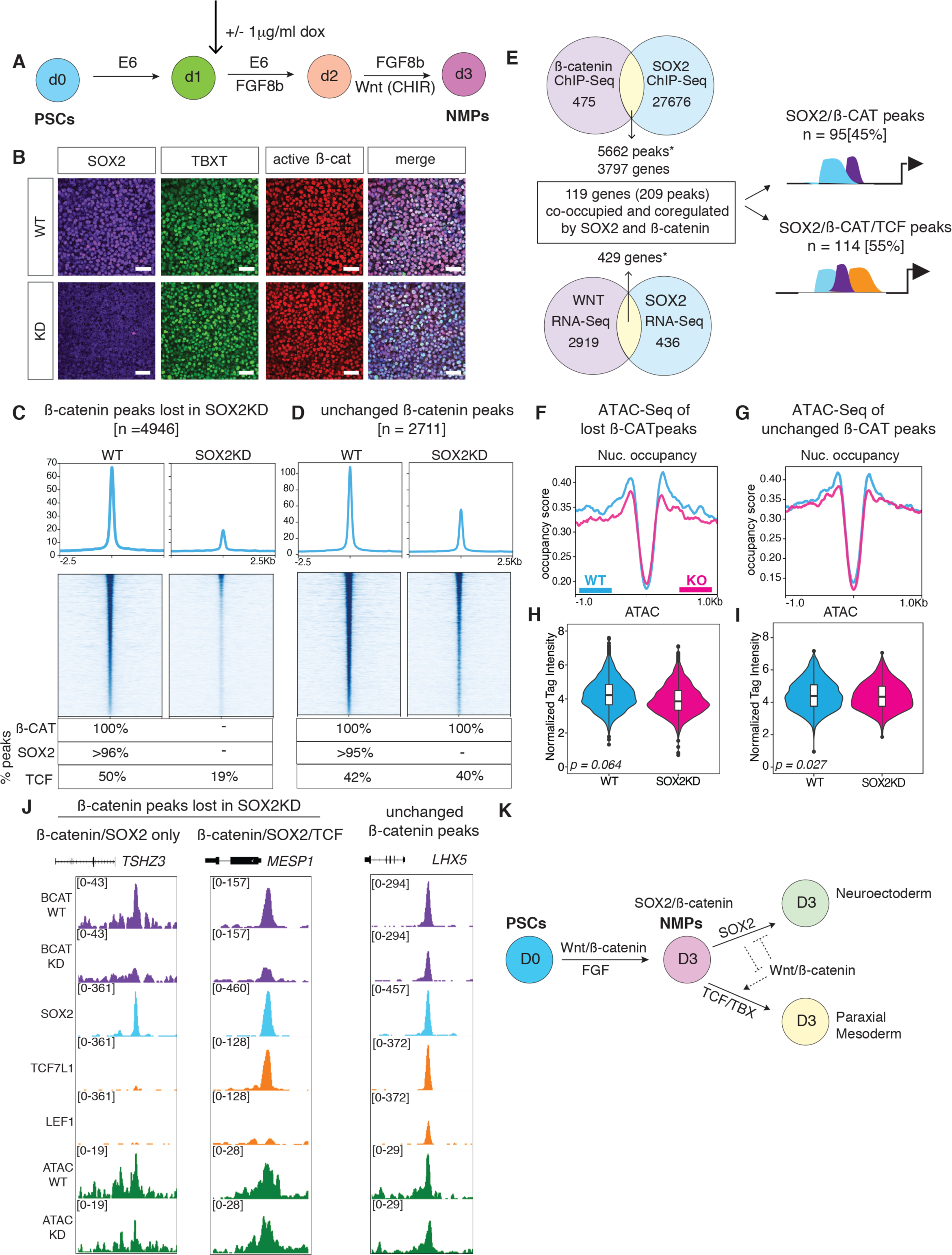
SOX2 is required for ß-catenin chromatin recruitment in neuromesodermal progenitors. **A.** schematic of neuromesodermal progenitor (NMPs) differentiation with SOX2KD CRISPRi induction by dox. **B.** Immunostaining for SOX2, TBXT and active ß-catenin protein levels in WT and SOX2KD cells on Day 3 of NMP differentiation. **C-D.** ß-catenin ChP-seq density plots and metaplots of the average signal intensity at ß-catenin peaks that are **C.** lost in SOX2KD cells and at **D.** peaks that remain unchanged in WT and SOX2KD cells (D). The table shows the percentage of peaks that were bound ß-catenin, SOX2 or TCF (either TCF7L1, LEF1 or both) for each category in WT or SOX2KD cells. **E.** Integration of ß-catenin and SOX2 RNA-Seq and ChIP-Seq datasets. *Significant overlap based on hypergeometric test, ß-catenin and SOX2 ChIP-Seq peak overlap: *p* 1.07 = × 10^-459^; WNT and SOX2 regulated genes sets from RNA-Seq: *p =* 5.64 × 10^-352^. **F-G.** ATAC-Seq nucleosome occupancy signal showing **F.** loci that lost ß-catenin peaks or **G.** ß-catenin peaks unchanged in SOX2KD. **H-I.** Violin plots quantifying ATAC-Seq read densities at **H**. loci that loose ß-catenin peaks in SOX2KD or **I.** loci where ß-catenin peaks were unchanged in SOX2KD. **J.** Representative genome browser views of loci that lostß-catenin or remain unchanged in WT and SOX2KD cells. **K.** Schematic summarizing SOX2 and WNT/ß-catenin interactions during NMP specification.

To characterize the genomic binding of SOX2 and in ß-catenin, we performed ChIP-Seq experiments in NMP cells. These experiments revealed that 92% [n = 5662/6137] of ß-catenin bound genomic loci are also occupied by SOX2 [Fig5E]. We then performed ß-catenin ChIP-Seq in WT and SOX2KD NMP cells [Fig S9A]. Differential peak analysis revealed that SOX2KD led to a significant loss of ß-catenin binding at 4946 loci, while ß-catenin binding was unchanged at 2711 loci and only 214 new ß-catenin peaks were gained [Fig 5C,D FigS9 A-E]. *De-novo* motif analysis revealed an enrichment of TCF DNA-binding sites in both lost and unchanged in ß-catenin peaks, while, as expected, SOX motifs were only enriched at loci that lost ß-catenin binding in SOX2KD cells [Fig S9F]. GO analyses of the genes associated with SOX2-dependent ß-catenin binding showed an enrichment for terms related to nervous system development, while unchanged ß-catenin peaks were enriched for terms related to mesoderm development. This is consistent with the idea that SOX2 promotes neural fate in NMPs while WNT favors mesoderm differentiation [Fig S9F].

To identify direct SOX2 and WNT target genes, we then performed RNA-seq on WT and SOX2KD cells as well as on cultures where CHIR was replaced with C59 to inhibit WNT signaling [Fig S8A, D; see Methods]. Differential expression analysis identified 865 SOX2 regulated and 2491 WNT regulated transcripts. GO analysis showed that the 346 genes downregulated in the SOX2KD were enriched for ‘epidermis development’ and neurogenesis’ whereas the 519 upregulated genes were enriched for terms related to ‘WNT signaling’ and ‘A-P axis specification’ [Fig S8B, C]. WNT regulated genes had a similar but opposite functional annotation with CHIR activated genes being enriched for ‘A/P axis specification’ and ‘mesoderm development’, while WNT repressed gene were enriched for terms associated with nervous system development [Fig S8E, F]. Integrating the ChIP-seq and RNA-seq data identified 209 enhancers, corresponding to 119 genes that were coordinately co-occupied and coregulated by SOX2 and ß-catenin [Fig 5E. S8I]. Consistent with an antagonistic relationship between SOX2 and WNT/ß-catenin, 42% of co-occupied and coregulated genes [50/119] were SOX2 repressed but WNT activated whereas 29% [34/119] were SOX2 activated and WNT repressed [Fig S8J].

Next, we evaluated TCF occupancy of the SOX2/ß-catenin co-bound loci by ChIP-seq for TCF7L1 and LEF1, the two TCFs most highly expressed in NMPs. We found that 50% [2483/4946] of the loci that lost ß-catenin peaks in SOX2KD cells were also co-occupied by TCFs like those associated with *MESP1*, while the other half had no evidence of TCF7L1 or LEF1 binding such as *TSHZ3* [Fig 5C-D, J; Fig S9 D – E.]. We then assessed chromatin accessibility at loci with SOX2-dependent ß-catenin binding by ATAC-Seq of WT and SOX2KD NMPs. Surprisingly, we did not observe appreciable differences in ATAC-seq signal at loci that had SOX2-dependent ß-catenin binding with the majority of these loci being accessible in both WT and SOX2KD cells [Fig S9G].

Collectively, our genomic analysis of SOX2/ß-catenin in NMPs and SOX17/ß-catenin in DE demonstrate that SOX TFs are required to regulate ß-catenin recruitment and lineage-specific WNT responsive transcription. In some cases, SOXs co-occupy WREs with TCFs, in other cases SOXs appear to recruit ß-catenin independent of TCFs.

### SOX17 assembles a WNT-responsive transcription complex at TCF-independent enhancers

To understand how SOX17 and ß-catenin activate transcription, we focused on a subset of SOX17/ß-catenin regulated TCF-independent enhancers. We tested a -60kb *CXCR4* and a - 33kb *BMP7* enhancer; both these genes were coregulated and co-occupied by SOX17 and ß-catenin, but had little evidence of TCF occupancy. DNA sequence analysis confirmed that both these putative enhancers had several SOX17 but no TCF binding sites [see Methods, Supplementary Table 4]. We termed this category of enhancers as ‘SOX-dependent’. Moreover, *CXCR4* is a well-established DE marker co-expressing with SOX17 in both the developing mouse endoderm and in human DE cultures^56, 57^. Interestingly, *BMP7* has been implicated as direct SOX17 target during germ-cell differentiation, supporting a similar relationship in DE^58^. As controls, we also assessed exemplar ‘universal’ WREs corresponding to *SP5*^59^ and *NKD1*^60^, that are activated through canonical ß-catenin/TCF interactions. We termed this category of enhancers as ‘TCF-dependent’. We cloned each of these enhancers into luciferase reporter constructs and also generated versions where the putative SOX17 DNA-binding sites or TCF sites were mutated (ΔSOX and ΔTCF respectively). We then transfected the WT or mutant enhancers into PSCs differentiated into WT DE, SOX17KO Day 3 cultures (-SOX17) or Day 3 cultures where ß-catenin activity was inhibited by removal of CHIR and addition of C59 (-WNT). The *CXCR4* and *BMP7* enhancers were both robustly active in WT DE and demonstrated significantly decreased activity upon addition of the C59 (-WNT) or in SOX17KO (-SOX17) cells. Moreover, mutation of the SOX17 DNA-binding sites dramatically reduced enhancer activity [Fig 6A].

**Figure 6.**
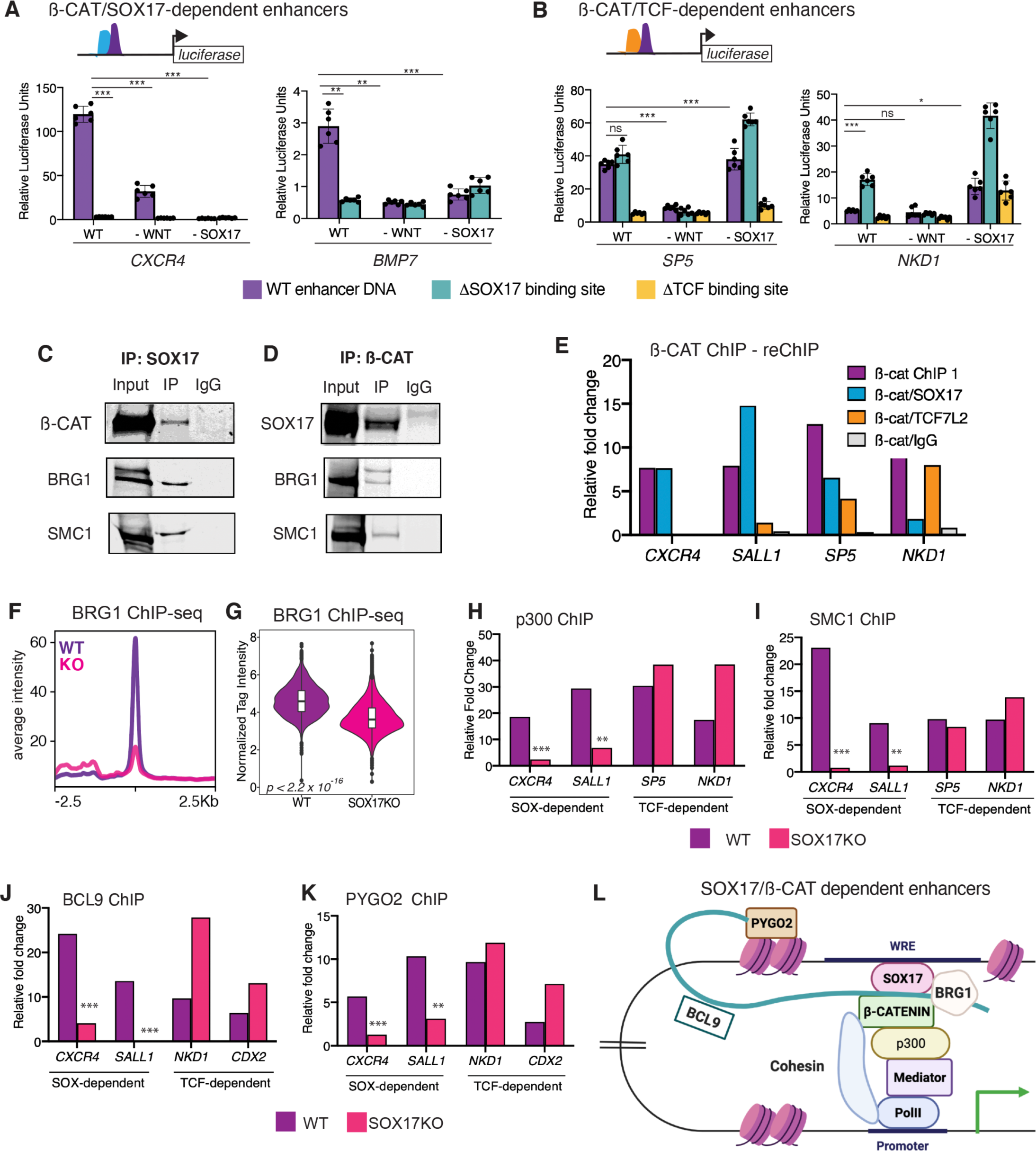
SOX17 can recruit a ß-catenin transcriptional complex to Wnt-responsive enhancers without TCFS. **A – B.** Schematic of enhancer luciferase constructs containing either wild type (WT) sequences or with SOX (blue) or TCF (orange) DNA-binding sites mutated. Histogram showing average luciferase reporter activity of **A.** SOX17-dependent or **B.** TCF-dependent enhancers in wild type (WT) cells, C59 treated Wnt inhibited cells (-WNT) or SOX17KO (-SOX17) cells. Statistical differences were determined between WT and ΔSOX17 enhancers in WT cells, WT enhancers in WT and -WNT cells, and WT enhancers in WT and -SOX17 cells by two-tailed student’s T-test. ns = not significant, * = *p<0.05,* **=*p<0.01,* ***=p<0.001. **C-D.** Western blots showing the presence of interacting partners followed by co-immunoprecipitation of **C.** SOX17 and **D.** ß-catenin. **E.** ChIP-reChiPIP-qPCR, showing the relative fold change in chromatin recovery with ß-catenin ChIP followed by a second reChIP with either SOX17, TCF7L2 or IgG at SOX-dependent (*CXCR4* and *SALL1*) or TCF-dependent (*SP5 and NKD1*) enhancers. **F-G** BRG1 ChIP-seq in WT (purple) and SOX17KO (pink) cells**. F.** Metaplots showing the average peak intensity at loci with SOX17-dependent ß-catenin binding which are not bound by TCFs. **G.** violin plots quantifying BRG1 read intensity of peaks in F. Wilcoxon rank sum test, p<2.2x10^-16^. **H – K.** ChIP-qPCR of p300 (H), SMC1 (I), BCL9 (J) and PYGO2 (K) showing relative fold change chromatin binding at SOX-dependent or TCF-dependent enhancers. Two-tailed student’s T-test. ns = not significant, * = *p<0.05,* **=*p<0.01,* ***p=<0.001. **L.** Model depicting SOX17-dependent assembly of a Wnt-responsive transcription complex at TCF-independent endodermal enhancers.

In WT DE, the *SP5* enhancer displayed robust activity, and this was not altered in SOX17KO cells. Motif analysis showed evidence of multiple TCF as well as SOX17 binding sites. Mutating the SOX17 sites did not affect enhancer activity, while as expected, mutating TCF sites led to a significant loss of reporter activity [Fig 6B]. This is consistent with the regulation of endogenous *SP5* [Fig. 1, 3]. In contrast, *NKD1* is an example of a gene activated by WNT but repressed by SOX17. Accordingly, the ‘wild-type’ *NKD1* enhancer had minimal transcriptional activity in WT DE cells, but was activated in SOX17KO cells, or by mutating SOX17 sites. On the other hand, mutating TCF sites led to decreased *NKD1* reporter activity [Fig 6B]. Together, these data demonstrate that TCF-independent SOX17/ß-catenin-bound loci are *bona fide* WNT-responsive enhancers. Further, our experiments recapitulate distinct modes of both SOX17 and TCF occupancy and target gene regulation at distinct subsets of WREs.

Next, we performed reciprocal coimmunoprecipitation (coIP) experiments demonstrating that SOX17 and ß-catenin physically interact in DE cells [Fig 6C, D]. As expected, we also observed a direct interaction between TCF7L2 and ß-catenin in both WT and SOX17KO cells [Fig S10A]. Reciprocal ChIP-reChIP experiments further confirmed that both SOX17 and ß-catenin directly interact at these SOX-dependent but not TCF-dependent enhancers [Fig 6E, FigS10 B].

We next tested the hypothesis that SOX-dependent WREs can serve as a scaffold for recruitment of transcriptional coactivators. ß-catenin interacts with chromatin modifiers through its C-terminal transactivation domain including BRG1 and p300^8, 61, 62^. ß-catenin also interacts with components of the cohesin and mediator complexes, previously shown to be critical for transactivation of TCF/ß-catenin target genes^63–65^. CoIP assays demonstrated that both ß-catenin and SOX17 interact with BRG1 and the cohesin subunit SMC1 in DE cells [Fig 6C, D]. To assess if SOX17 was required to recruit BRG1 to TCF-independent enhancers, we performed BRG1 ChIP-Seq in WT and SOX17KO cells. This revealed a substantial loss of BRG1 signal in SOX17KO cells at those TCF-independent enhancers that we previously demonstrated exhibited SOX17-dependent ß-catenin binding [Fig 4, 6F, G]. We then performed ChIP-qPCR assays in WT and SOX17KO cells for other previously known interactors of TCF/ß-catenin transcriptional activation complexes including p300^66^, the cohesin subunit SMC1, the cohesin loading protein NIPBL^67^ the mediator subunit MED12, as well as the core WNT enhanceosome components BCL9 and PYGO^68–70^ [Fig 6H – K, FigS10 D-E]. In each case, SOX17 was required for efficient recruitment to the TCF-independent SOX17/ß-catenin regulated DE enhancers *CXCR4* and *SALL1*. In contrast, there was no difference in occupancy of these coactivators to the TCF-dependent WREs *SP5*, *NKD1* and *CDX2* in SOX17KO cells [Fig. 6H-K, Fig S10D-E]. We did not detect any differences in expression levels of WNT enhanceosome components or transcriptional coactivators in WT and SOX17KO cells, suggesting that SOX17 is directly required to recruit these interacting partners to SOX-dependent enhancers [Fig S10C].

Collectively, our data suggests that SOX17 is required to recruit ß-catenin to a subset of DE enhancers and assemble a TCF-independent transcription complex to activate lineage-specific WNT-responsive transcription [Fig 6L].

## DISCUSSION

### Overview

In this study, we tested the hypothesis that the SOX family of TFs function as lineage-specific regulators of WNT responsive transcription. We showed that during hPSC-derived DE differentiation, the recruitment of ß-catenin to lineage-specific enhancers was highly dynamic and cannot be accounted for completely by TCFs. During DE differentiation, there was an increased number of genomic loci co-occupied by ß-catenin and SOX17. The loss of SOX17 led to widespread genomic relocalization of ß-catenin binding, with SOX17 being required for the recruitment ß-catenin to a subset of enhancers that are WNT-responsive and SOX17-regulated, but have no evidence or TCF binding. At some of these TCF-independent enhancers, SOX17 and ß-catenin interacted with Wnt-pathway components BCL9 and PYGO as well as transcriptional coactivators p300, BRG1, MED12 and SMC1 to assemble a SOX17-dependent transcription complex. Mutating SOX17 DNA binding sites led to a loss of transcriptional activity of these enhancers. Similarly, we showed that SOX2 is also required for the chromatin recruitment of ß-catenin to regulate lineage-specific Wnt-responsive transcription in hPSC-derived NMP cells; half of these loci also had no evidence of TCF co-occupancy. Although we have focused on the most novel cases where SOXs appear to recruit ß-catenin independent of TCFs, in both DE and NMPs, many genomic loci are also co-occupied by SOXs, TCFs and ß-catenin. Moreover, we observe many WNT-responsive genomic loci that retain ß-catenin binding irrespective of SOX17 depletion, as well as a substantial proportion of loci that gain *de-novo* TCF/ß-catenin binding upon loss of SOX17. These data suggest that the interplay between SOXs and TCFs is likely to be more complex and that both cooperative and competitive interactions between SOX and TCF TFs may regulate ß-catenin recruitment to distinct context-specific loci.

### TCF-independent WNT-responsive transcription

According to the current dogma, ß-catenin/TCF interactions mediate the vast majority, if not all, of WNT responsive transcription. Indeed, in HEK293T and intestinal epithelial cells, ß-catenin binding events almost completely overlap with TCFs, and dominant negative TCF7L2 is sufficient to diminish ß-catenin binding at the vast majority of peaks^71^. While the central role of TCFs in mediating Wnt-responsive transcription is not in doubt, it is still unclear whether TCFs can account for the full diversity of Wnt-responsive transcription in different cell types, particularly during development. Indeed, a recent study showed that despite deletion of all four TCFs in HEK293T, ß-catenin retained transcriptional activity and binding at some genomic loci.^72^ While other TFs have anecdotally been shown to bind to ß-catenin in various contexts (often *in vitro*) the notion that alternative TFs had a major role in mediating WNT-responsive transcription, independent of TCFs has been controversial.

Our systematic time-resolved genomic analyses shows that TCFs cannot account for the full extent of ß-catenin chromatin binding during DE differentiation. While ß-catenin binding and Wnt-responsive transcription is almost exclusively mediated by TCFs in Day 1 mesendoderm cells, SOX17 accounts for an increasing proportion of ß-catenin binding event as DE differentiation progresses. Like SOX17, SOX2 can also regulate lineage-specific WNT responsive transcription by directing ß-catenin recruitment to lineage-specific neuromesodermal and neuronal loci. Collectively, our studies indicate that SOX TFs account for a large proportion of the genomic ß-catenin binding in two different developmental lineages. One possibility is that ß-catenin/TCF interactions preferentially regulate context-independent functions, such as WNT-mediated cell proliferation. In contrast, TCF-independent ß-catenin/SOX or ß-catenin/SOX/TCF interactions may be more prevalent during development where transcriptional programs are more dynamic. Further studies discriminating between these scenarios shall provide more insight into how specific ß-catenin is preferentially engaged by TCF vs. non-TCF TFs.

### SOX TFs as lineage-specific regulators of the WNT pathway

Most, if not all SOX TFs are reported to modulate WNT responsive transcription through reporter assays (TOP:flash) in overexpression conditions^28^. Similar to the TCFs, the SOX TFs bind the minor groove of DNA through their HMG domains and induce a bend of 60-70°, facilitating interactions with local chromatin modifiers and transcriptional coactivators and leading to transcription initiation^73^. Our study highlights the role of SOX17 and SOX2 as dual-function regulators of endodermal and neuromesodermal lineages, respectively: not only are they required to activate lineage-specific GRNs, but they directly repress alternate-lineage fates.

Despite binding DNA through low affinity sequences, the SOX TFs demonstrate remarkable specificity in gene regulation. One attractive consideration is that they form lineage-specific regulatory complexes with homologous or heterologous partners, thereby providing specificity towards the regulation of its target genes. A classic example of the SOX ‘partner-code’^74^ is the SOX2-OCT4 heterodimer that’s critical for maintenance of pluripotency in PSCs^24^, and in mouse ESCs, they can physically associate with ß-catenin/Tcf7l1^75^. SOX TFs also display remarkable differences outside of the HMG domain^23^. Accordingly, SOX17, a SOXF group member, cannot substitute for the functions of SOX2, a member of the SOXB1 group. Mutating an acidic glutamate residue within SOX17 to lysine, however, enables it to bind to OCT, and subsequently SOX2 binding sites at pluripotency related loci^76, 77^. In the future, it would be interesting to identify and test if SOX subgroup-specific domains outside of the HMG-box are required for selective ß-catenin binding to lineage-specific loci.

Our ChIP-seq analyses show a large degree of overlap between ß-catenin, SOX17 and GATA6/GATA4 binding events during DE differentiation. This is consistent with previous studies in *Xenopus* embryos where Sox17 and Gata4/6 coordinately regulate endoderm development^27, 78^, as well as recent studies that GATA6 functions upstream of SOX17 and FOXA2 to pattern an endoderm-specific chromatin landscape and can directly interact with SOX17^79^. Interestingly, ß-catenin/TCF7L2/GATA4 and TCF7L2/GATA2/GATA1 interactions have also been reported during cardiac and erythroid lineages specification respectively^80, 81^. It is possible that SOX17-GATA heterodimers recruit ß-catenin to DE-specific enhancers. Future computational analyses and biochemical experiments will allow us to distinguish between monomeric and multimeric SOX17 sites and whether ß-catenin is preferentially recruited solely by SOX17 or by a cluster of lineage-specific TFs.

### Multiple modes of interactions between SOX and TCF TFs

Our studies reveal multiple modes of interactions between SOX and TCF TFs. In addition to TCF-independent enhancers, we identify a subset of enhancers co-occupied by SOX17 and TCFs where ß-catenin occupancy remains unchanged in the presence and absence of SOX17. This is consistent with previous studies showing that lineage-directing TFs like SOX and signal-determining TFs like TCFs act combinatorically on tissue-specific enhancers; this is one likely mechanism directing lineage-specific Wnt-responsive transcription at a subset of genomic loci^20, 81^. *In-vitro* protein binding assays using recombinant Sox17, Tcf7l2 and ß-catenin have shown that they can form a trimeric complex^26^. Interestingly, at endogenous levels in DE cells, we could not detect a physical interaction between SOX17 and TCF7L2. Sequence analysis of enhancers co-occupied by SOX17 and TCFs show that the majority of SOX17 and TCF binding sites are >50bp away from each other. However, since they are both HMG box TFs that bend DNA, it is conceivable that despite being distant from each other on linear DNA, they might cooperatively bind ß-catenin. On the other hand, our results show that the global increase in *de-novo* ß-catenin/TCF binding in SOX17KO cells. These points to an interesting possibility that SOX17 and TCFs compete for recruitment of a finite amount of nuclear ß-catenin. Future genomic and biochemical experiments with careful titration of SOX and TCF levels will be important to test this.

### The WNT/ß-catenin enhanceosome complex

Several studies have recently identified nuclear proteins required for the assembly of a ß-catenin-responsive transcription complex – termed the WNT enhanceosome^9, 10^. While components such as PYGO potentiate the transactivation of WNT target genes, BCL9 acts as a bridge to tether ß-catenin to PYGO^68, 82^. It’s been proposed that upon WNT stimulation and recruitment of ß-catenin to chromatin by TCFs, the WNT enhanceosome undergoes a conformational change to bring the distal enhancer in close proximity to its cognate promoter, leading to the recruitment of RNA PolII and transcriptional activation^67^.

The notion that TFs other than TCF might function to integrate the multiple components of this enhanceosome has been proposed previously^11^. For example, TBX3 associates with the WNT enhanceosome through interactions with BCL9^22^ and the RUNX family of TFs can interact with the ChiLS complex^9^. However, the extent to which these TFs can regulate ß-catenin genomic recruitment is unknown.

Our experiments show that on TCF-independent WNT responsive enhancers, SOX17 is required for the recruitment of BCL9, PYGO2, as well as transcriptional coactivators p300, BRG1, MED12 and SMC2, which physically interact with the ß-catenin C-terminal transactivation domain. However, the hierarchy and the sequential order in which SOX17 and the WNT enhanceosome complex are assembled on DNA remains to be determined. An attractive hypothesis supported by our data is that SOX17 acts in a sequential manner to first increase the chromatin accessibility at specific enhancers, perhaps acting as a pioneer TF. This then sets the scene for SOX17 to recruit enhanceosome components like Pygopus which facilitates the loading ß-catenin, similar to a model that’s been proposed to prime ß-catenin/TCF enhancers to respond to WNT activation^9^. Ultimately, elucidation of the SOX17/ ß-catenin/DNA complex structure, coupled with proteomics and biochemical assays will be critical to dissect the mechanisms assembling a SOX17/ß-catenin transcription complex.

Another interesting possibility, in the context of enhancers cobound by SOX, ß-catenin and TCF, is that SOX TFs might recruit coactivators or corepressors to potentiate or repress traditional TCF/ß-catenin. Indeed, SOX9 is reported to enhance ß-catenin phosphorylation and turnover in chondrocytes^31, 83^, and in SW480 cells, SOX17 negatively regulates WNT responses^26^. While our results show no appreciable differences in ß-catenin or TCF protein levels between WT and SOX17KO cells, it is possible that SOX17 instead regulates the WNT enhanceosome through multiple mechanisms including perhaps post-translational modifications of ß-catenin, such as trimethylation or acetylation, affecting activation vs. repression^37^, or via chromatin modifiers such as *Kdm2a/b* to regulate stability of nuclear ß-catenin at specific loci^84^. Further studies are needed to explore if and how SOX TFs recruit proteins that post translationally modify ß-catenin at specific loci.

In summary, the data here establish the SOX TFs as context-dependent regulators of WNT-responsive transcription by regulating the recruitment of ß-catenin and enhanceosome complexes to lineage-specific enhancers. Given that most, if not all cell types, express at least one of the 20 SOX TFs that are encoded in the human genome, it is likely that they have a broad, previously unappreciated role in regulating WNT responsive transcription in many contexts.

Indeed, there is correlative evidence of SOX TFs and ß-catenin interactions leading to the dysregulation of WNT-responsive oncogenes in many cancers, including breast, cervical and colon cancers^30, 85, 86^. Therefore, further investigations into the mechanisms through which SOX and TCFs interact to control the genomic specificity of ß-catenin in different cellular contexts might open up the possibility of targeting ß-catenin-SOX interactions for therapeutic purposes.

### DATA AVAILABILITY

Datasets generated in this study have been deposited to the Gene Expression Omnibus (GEO): GSE # pending. A description of all datasets generated in this study can be found in Supplementary Table 1.

## Supporting information

Datasets_Metadata

Key_Reagents

FigS1_Stats

## ACKNOWLEDGEMENTS

We thank Keely Icardi and the CCHMC DNA Sequencing Core for help with ChIP-Seq library preparation; CCHMC Pluripotent Stem Cell Facility and Evan Brooks for help with generating the SOX17KO line. The CRISPRi-SOX2 line was a kind gift from Dr. Bruce Conklin (Gladstone Institutes, UCSF). This work was supported by NIDDK R01 DK123092 and in part by NIH P30 DK078392 to A.M.Z.

## AUTHOR CONTRIBUTIONS

S.M. and A.M.Z. designed the project, interpreted data and wrote the paper. S.M. performed all experiments and data analyses in collaboration with: D.M.L for cDNA cloning, immunofluorescence, and western blot experiments and L.B. for reporter assays. All authors provided input and approved of the final version of the paper.

## METHODS

### Cell Culture

Human embryonic stem cell line WA01 (H1) was purchased from WiCell and induced pluripotent stem cell line iPS72.3 was obtained from CCHMC Pluripotent Stem Cell Facility. The CRISPRi-SOX2 line was a kind gift from Dr. Bruce Conklin (Gladstones Institutes, UCSF). hESCs and iPSCs were maintained in feeder-free cultures. Cells were plated on hESC-qualified Matrigel (Corning; 354277) and maintained on mTESR1 (StemCell Technologies; 85851) media at 37°C with 5% CO2. Media was changed daily, and cells were routinely passaged every 4 days using ReleSR (StemCell Technologies; 05872). CRISPRi-SOX2 cells were plated on vitronectin-coated (ThermoFisher; A14700) plates and maintained in Essential 8 Medium(Gibco; A1517001). These lines were routinely passaged every 3-4 days using Versene (ThermoFisher; 15040066). All lines were routinely screened for differentiation and tested for mycoplasma contamination.

### Generation of SOX17KO line

*<u>gRNA Validation:</u>* Two CRISPR/Cas9 guide RNAs targeting the first exon of the SOX17 gene were cloned into pX458 (Addgene 48138) and validated in HEK293T cells (ATCC CRL-3216) by the Transgenic Animal and Gene Editing core at CCHMC. The CRISPR targeted region was amplified with Phusion Polymerase (ThermoFisher; F531) and each amplicon was digested with T7 Endonuclease I (NEB; M0302S). Digested amplicons were run on agarose gels to quantify relative gRNA activity. *<u>Nucleofection:</u>* RNP complex assembly of the validated gRNA was performed by combining 20ug Alt-R® S.p. HiFi Cas9 Nuclease V3 (IDT; 1081060) with 16ug sgRNA (Synthego) *in vitro* for 45 minutes at room temperature. The RNP complex was electroporated into parental iPS 72.3 cells using a Lonza 4D Nucleofector. Isolated clones were lysed and amplified using Phusion polymerase and clones of interest were submitted for Sanger sequencing to the CCHMC DNA Sequencing Core.

### Definitive Endoderm differentiation

Confluent cells were passaged to single cells using Accutase (Sigma Aldrich; A6964) and plated on Matrigel-coated plates using mTESR1 and Y-27632 (StemCell Technologies, 72304). The following data, basal media was replaced, and cells were washed with PBS. DE differentiation was then carried out in RPMI-1640 media (Thermo Fisher; 11875-093) supplemented with non-essential amino acids (ThermoFisher; 11140050). Cells were treated with 100ng/ml Activin A (Shenandoah, 800-01) and 2um CHIR99021 (R&D Systems; 4423) for 24hrs. In the next two days, cells were treated with 100ng/ml Activin A and 2μm CHIR99021 in RPMI-1640 with increasing concentrations (0.2% on Day 2, 2% on Day 3) of ES-grade FBS (GE; SH30070.02). To identify Wnt-responsive genes, cells were treated with 1μm of the Wnt inhibitor C59 (Tocris; 5148) at two different time points; for 3 days from the onset of differentiation (between Day 0 and Day 1) to identify ‘early’ Wnt regulated genes and on days 2 and 3 to identify ‘late’ Wnt regulated genes.

### Neuromesodermal Progenitor differentiation

Confluent cells were passaged using Accutase and plated on vitronectin coated plates using E8 and Y-27632. NMP differentiation was carried out largely as described previously^55^. Briefly, media was changed to Essential 6 Medium (Gibco; A1516401), 24 hours later cells were treated with 200ng/ml FGF8b (PeproTech; 100-25) in E6 media. After a further 24 hours, cells were treated with 200ng/ml FGF8b and 3μm CHIR99021 in E6 media. To knock down SOX2 levels, the CRISPRi-SOX2 cells were treated with 1μg/ml doxycycline (dox) on days 2 and 3 of differentiation. To identify Wnt regulated genes, NMP cultures were treated with either 3μm CHIR99021 or 1μm C59 on Day 3 of differentiation.

### mRNA Extraction. RT-qPCR and RNA-Seq

Total RNA was extracted using the Nucleospin RNA Extraction kit (Machery-Nagel; 740955) and reverse-transcribed to cDNA using SuperScript VILO (ThermoFisher; 1177250) according to manufacturer’s instructions. qPCR was performed using PowerUp SYBR Green MasterMix (ThermoFisher; A25777) and QuantStudio 3 Flex Real-Time PCR system. Relative mRNA expression was normalized to that of housekeeping gene *PPIA (*peptidylprolyl isomerase A) and calculated using the *ΔΔ*Ct method. For RNA-Seq experiments, three biological replicates were sequenced per condition. 300ng of total RNA, as determined by Qubit High-Sensitivity spectrofluorometric measurement, was poly-A selected and reverse transcribed using Illumina’s TruSeq stranded mRNA library preparation kit (Illumina; 20020595). Samples were incubated with unique Illumina-compatible adapters for multiplexing. After 15 cycles of amplification, libraries were paired end sequenced on a NovaSeq 6000 with a 2x100 read length.

### Immunofluorescence

Cells were plated at a density of 10,000 cells/ml on Matrigel or vitronectin coated Ibidi 8-well chamber slides (Ibidi; 80826). Cells were washed once with PBS and fixed in 4% paraformaldehyde for 30 minutes at room temperature. If necessary, antigen retrieval was performed by adding 1x Citrate Buffer warmed to 55°C and incubating slides at 65°C for 45 mins.

Slides were then blocked with 5% normal donkey serum (NDS) for an hour. Primary antibodies were added in 5% NDS in PBS and incubated overnight at 4°C. The following day, cells were washed thrice in PBS and incubated with secondary antibodies and DAPI for an hour at room temperature. Slides were again washed in PBS before imaging. Images were taken using a Nikon A1R inverted confocal microscope and analyzed using NIS Elements (Nikon). Antibodies and dilutions used are listed in Supplementary Table 2.

### Cell Fractionation

Nuclear isolation was performed as previously described^87^. Briefly, cells were dissociated using Accutase and counted using a Bio Rad TC20 Automated Cell Counter. 10 million cells were then lysed in 1ml of cytoplasmic buffer (50mM Tris-HCl pH 7.5, 10% glycerol, 0.5% Triton X-100, 137.5mM NaCl) supplemented with protease (ThermoFisher; A32953) and phosphatase (ThermoFisher; A32957) inhibitors and incubated on ice for 15min. Cells were then pelleted by centrifugation for 5 mins at 16,000 rpm at 4°C. Nuclei were then resuspended in 10mM HEPES pH 7.8, 0.5 M NaCl, 0.1% NP-40 supplemented with 1mM DTT and fresh protease/phosphatase inhibitors. Samples were sonicated for two 10s pulses on ice. Nuclei were cleared by centrifugation for 10mins at 16,000 rpm at 4°C.

### Co-immunoprecipitation

CoIP assays were performed as previously described^88^ with minor modifications. After differentiation to the desired stage, 20 million cells were washed on the plates with PBS and crosslinked with 1.5mM DSP (ThermoFisher; 22585) for 30 mins at room temperature. The crosslinking reaction was quenched by 30mM Tris pH 7.4 in PBS and incubated for 20 minutes. Cells were then scraped in ice-cold PBS supplemented with protease inhibitors and resuspended in 1ml cytoplasmic buffer (10mM HEPES pH 7.9, 10mM KCl, 340mM sucrose, 3mM MgCl2, 10% glycerol, 0.1% TritonX-100) supplemented with 1mM DTT and fresh protease inhibitors. Cells were incubated on ice for 10 mins and centrifuged for 10 mins at 4000 rpm. The nuclear pellet was then resuspended in 500μl CoIP wash buffer (100mM NaCl, 25mM HEPES pH 7.9, 1mM MgCl2, 0.2mM EDTA, 0.5% NP-40) supplemented with protease inhibitors. Samples were sonicated for two 10s pulses, treated with 600U/ml benzonase (Millipore; 70664) and incubated at 4°C with end over end rotation for 4 hrs. Afterwards, the concentration of NaCl in the samples was adjusted to 200mM and samples were incubated for an additional 30 mins. Nuclear extracts were then cleared by centrifuging for 30 minutes at max speed at 4°C. Nuclear lysates were then quantified by BCA assays and protein concentrations of the lysates were adjusted to either 500ug/ml or 1mg/ml by diluting with CoIP wash buffer. Lysates were then precleared with Protein G Dynabeads (ThermoFisher; 10004D) at 4°C for an hour. 10% input samples were collected from the precleared lysates and stored at -20°C. Then samples were transferred to fresh tubes and incubated with relevant antibodies overnight at 4°C with end-over-end rotation. The following day, lysates and antibody complexes were added to precleared Protein-G Dynabeads and allowed to incubate at 4°C for 2 hrs with end-over-end rotation. The antibody/beads complexes were then washed with ice-cold CoIP wash buffer 8 times at 4°C. Lysates were then briefly centrifuged to remove any residual wash buffer and the beads were resuspended in 60μl 2x LDS loading buffer (ThermoFisher; NP0007). Proteins were eluted from beads on a thermomixer at 65°C for 15 mins at 1000 rpm. Immunoprecipitations with antibody and IgG were performed parallelly. A list of antibodies and associated dilutions can be found in Supplementary Table 2.

### Western Blots

Nuclear lysates were quantified by BCA and equal concentrations of protein samples were loaded for all experiments. Samples were resuspended in 4x LDS loading buffer supplemented with fresh 100mM DTT and boiled for 10mins. Proteins were separated on 4-12% Bis-Tris or 7% Tris-Acetate gels and transferred to PVDF membranes. Membranes were blocked in LI-COR TBS Intercept Blocking Buffer (LiCor; 927-60001) for an hour and then incubated with primary antibodies overnight at 4°C. The next day, membranes were probed with relevant secondary antibodies and imaged on a LI-COR Odyssey Clx scanner and processed using LI-COR Image Studio Lite. A list of antibodies and associated dilutions can be found in Supplementary Table 2.

### Transfections and Reporter Assays

Putative SOX17-dependnent or TCF-dependent enhancers were synthesized (Genscript) and cloned into the pGL4.23 (*luc2*/miniP; Promega) vector. For transfections, hESCs were dissociated into 2-3 cell clumps using Versene and plated at a density of 60,000 cells/ml using mTESR and RevitaCell supplement (Gibco; A2644501). DE differentiations were carried out as described above. On the completion of day 2 of differentiation, cells were washed with PBS and supplemented with fresh day 3 differentiation media and incubated at 37°C for 30 mins. 50μl Opti-MEM (Gibco; 31985062), 1μl Lipofectamine STEM Transfection Reagent (Invitrogen; STEM00001) and 500 ng DNA (495ng enhancer/luc, 5ng Renilla) were then added to the cells and they were incubated for 24hrs at 37°C. The next day, cells were washed with PBS, lysed and assayed using the Dual-Luciferase Assay System (Promega; E1910) according to manufacturer’s instructions.

### ChIP-qPCR, ChIP-reChIP and ChIP-Seq

Most ChIP experiments were performed in biological duplicates as in^71^ with several modifications. After differentiation to the desired stage, approximately 20 million cells were dual crosslinked in plate, first with 1.5mM EGS (ThermoFisher; 21565) for 20 mins, followed by supplementation with 1% formaldehyde for an additional 20 minutes at room temperature. The crosslinking reaction was quenched with 125mM glycine for 15 minutes at room temperature. Cells were then washed twice and scraped in ice-cold PBS and if needed, flash frozen in dry ice until future use. ChIP samples were resuspended in 1ml sonication buffer (20mM HEPES pH 7.4, 150mM NaCl, 0.1% SDS, 1% Triton X-100, 1mM EDTA, 0.5mM EGTA) supplemented with fresh protease inhibitors. Chromatin was sonicated using a Diagenode Bioruptor Pico instrument for 45 cycles of 30 seconds ON, 60 seconds OFF, to generate 200-400 bp sheared fragments. Chromatin was then precleared with Protein G Dynabeads for an hour with end-over-end rotation at 4°C. A volume of the precleared chromatin corresponding to 1% of the total volume was set aside as input. The rest of the samples were transferred to fresh tubes containing preblocked Protein G Dynabeads; relevant antibodies (see Supplementary Table 2) were added and samples were incubated overnight at 4°C with end-over-end rotation. The next day, the beads were washed serially with 150mM salt wash buffer (20mM HEPES pH 7.4, 150mM NaCl, 0.1% SDS, 0.1% sodium deoxycholate, 1% Triton X-100, 1mM EDTA, 0.5 mM EGTA), 500mM salt wash buffer (20mM HEPES pH 7.4, 500mM NaCl, 0.1% SDS, 0.1% sodium deoxycholate, 1% Triton X-100, 1mM EDTA, 0.5 mM EGTA), 1M salt wash buffer (20mM HEPES pH 7.4, 1M NaCl, 0.1% SDS, 0.1% sodium deoxycholate, 1% Triton X-100, 1mM EDTA, 0.5 mM EGTA), 2M salt wash buffer (20mM HEPES pH 7.4, 2M NaCl, 0.1% SDS, 0.1% sodium deoxycholate, 1% Triton X-100, 1mM EDTA, 0.5 mM EGTA) and LiCl wash buffer (20mM HEPES pH 7.4, 0.5M LiCl, 0.5% NP-40, 0.5% sodium deoxycholate, 1mM EDTA, 0.5 mM EGTA). Each wash buffer was supplemented with fresh 1mM DTT and protease inhibitors, and each wash was performed for 20 minutes at 4°C. The beads were then washed twice in 1xTE buffer supplemented with fresh protease inhibitors. The beads were then resuspended in DNA elution buffer (1% SDS, 0.1M NaHCO3). Elution was performed twice on a thermomixer at 65°C at 1,200 rpm. Eluates from two rounds of elution were combined and supplemented with 1/10 volume of 5M NaCl. Simultaneously, an equal volume of DNA elution buffer and 5M NaCl were added to the input samples. All samples were reverse crosslinked overnight at 65°C. The following day, samples were treated with RNase A for an hour at 37°C and digested with Proteinase K for 2 hours at 55°C. DNA was then purified using the Qiagen QIAquick Purification Kit (Qiagen; 28104) using manufacturer’s instructions and eluted in 20μl Elution Buffer; 1μl was used to quantify DNA concentrations using a Qubit High-Sensitivity DS DNA Assay kit (Invitrogen; Q32851).

For ChIP-Seq experiments, DNA libraries were prepared using 1-5 ng of starting material using the SMARTer ThruPLEX DNA-Seq kit (Takara; R400674) according to manufacturer’s instructions. After library amplification, DNA was purified using AMPure XP beads (Beckman-Coulter; A63880) and size-selected to retain 200-600 bp fragments. DNA fragment traces were analyzed on a Bioanalyzer. Suitable libraries were then paired-end sequenced on a Illumina NovaSeq 6000 with a 2x75 read length.

For ChIP-reChIP experiments, before the first ChIP, antibodies were crosslinked to Protein G Dynabeads. Briefly, beads were washed with 0.2M sodium borate pH9, and antibodies were crosslinked to beads by using 20mM DMP (Pierce; 21666) dissolved in 0.2M sodium borate. The crosslinking reaction was carried out at room temperature for 40 mins. The reaction was then quenched using 0.2M ethanolamine pH 8.0 for an hour. Residual IgGs were removed by washing the antibody/beads complex with 0.58%v/v acetic acid + 150mM NaCl. The beads were then added to processed chromatin samples and ChIP experiments were performed as described above. After the first ChIP, samples were eluted in DNA elution buffer supplemented with 10mM DTT. The samples were then diluted in 10 volumes of sonication buffer and the 2^nd^ ChIP was carried out as described above. qPCR was performed using PowerUp SYBR Green MasterMix and the QuantStudio 3 Flex Real-Time PCR system using default protocols. Primers were designed to span relevant SOX17 or ß-catenin peak centers and relative expression was normalized to that of a ‘negative’ control gene desert genomic region. Relative fold change was calculated using the *ΔΔ*Ct method. Primer sequences are listed in Supplementary Table 2.

### ATAC-Seq

ATAC-Seq experiments were largely performed as previously desribed^89^. Briefly, 50,000 cells were collected following differentiation to the desired stage, and lysed in 50μl of ATAC-lysis buffer (10mM Tris-HCl, pH 7.4. 10mM NaCl, 3mM MgCl2, 0.1% NP-40) to obtain a crude nuclei prep. All centrifugation steps were performed at 4°C at 2000 rpm. The nuclei pellet was then resuspended in the 50ul of the transposition reaction mix (25ul Tagment DNA buffer, 2.5ul TD Tn5 Transposase enzyme, 22.5ul nuclease-free water) (Illumina; 20034197). The transposition reaction was incubated at 37°C for 30 min on a thermomixer with constant gentle shaking at 1000 rpm. Immediately after transposition, DNA was purified using a Qiagen MinElute PCR Purification (Qiagen; 28004 kit) and eluted in 10μl Elution Buffer. The eluted DNA was then amplified in a reaction with 25μl NEBNext High-Fidelity 2x PCR Mastermix (NEB; M0541L) and custom 25um Nextera PCR Primers (Ad1_noMx universal primer, 0.5um Ad2.x indexing primer). PCR was performed as follows: 1 cycle of 72°C for 5min, 98°C for 30s, 5 cycles of 98°C for 10s, 63°C for 30s, 72°C for 1min. 5ul of the amplified DNA was then used to perform qPCR to determine the optimal number of additional cycles to prevent amplification saturation of DNA libraries. In all cases, either 4 or 5 additional cycles of PCR was performed at: 98°C for 10s, 63°C for 30s and 72°C for 1 min. Double size selection of amplified libraries (0.5x – left sided, 1.8x – right sided) was performed using AMPure XP beads and the DNA was eluted in a final volume of 20ul in 0.1x TE buffer. Purified libraries were sequenced on an Illumina NextSeq 500 instrument with a 2x75bp read length.

### Statistics and Reproducibility

RNA-Seq experiments were performed in biological triplicates. Most ChIP-Seq and ATAC-Seq experiments were performed in biological duplicates except for H3K27ac and BRG1 ChIP-Seq (n = 1) and NMP ß-catenin ChIP-Seq (n = 3). A description of all datasets generated in this study can be found in Supplementary Table 1. Immunostaining and western blots were performed at least four times and representative images were used. CoIP, ChIP-qPCR experiments and reporter assays were repeated at least thrice. All differentiations and experiments were performed using cell lines maintained between passages 55 – 65. To validate that CHIR-dependent target genes were indeed Wnt regulated, we also performed differentiation of iPSCs replacing CHIR99021 with recombinant WNT3A (R&D; 5036-WN) and validated the expression of several target genes by qPCR and ß-catenin and SOX17 binding by ChIP-qPCR.

## Data Analysis

### RNA-Seq

Raw reads were quality-checked using FASTQC (https://www.bioinformatics.babraham.ac.uk/projects/fastqc/) and if necessary, adapters were trimmed using cutadapt^90^. Fastq files were pseudo-aligned to the hg19 reference index using salmon^91^. An index of transcripts was built using default parameters (salmon index) using quasi-mapping (-quasi) and kmers of length 31 (-k 31). Relative transcript abundance was then quantified using salmon -quant using paired end fastq files and counts per transcript were obtained. The tximport package ^92^was then used for downstream analysis to convert transcript-level counts to gene-level estimates. Differential gene expression analysis was performed using DESeq2^93^ using default parameters. Differentially expressed genes were defined as those with log2 fold change >|1| and adjusted FDR of p<0.05. Transcripts were then annotated using the biomaRt ^94^ package. Any genes with TPM less than 10 across all replicates were then discarded from further analysis. To perform principal component analysis, variance stabilized and transformed data from DESeq2 *vst* function was generated. PCA plots were visualized using *plotPCA* () function of DESeq2.

To identify endoderm, mesoderm or ectoderm enriched genes, we reanalyzed the following datasets: GSM1112846, GSM1112844 (RNA-Seq of Day 3 ectoderm cells) and GSM1112835, GSM1112833 (RNA-Seq of Day 3 mesoderm cells) and compared with our Day 3 endoderm RNA-Seq data. Raw RNA-Seq data was downloaded from GEO and processed as described above, and pairwise differential gene expression analysis was performed using DESeq2 to identify genes with enriched expression in Day 3 endoderm, mesoderm or ectoderm. For instance, a gene was considered to be significantly endoderm enriched if: the gene was significantly differentially expressed in endoderm over control pluripotent cells, and showed significantly enriched expression in the endoderm compared to the mesoderm and ectoderm datasets. Differential enrichment threshold: log2 fold change >|1| and adjusted FDR of p<0.05.

### ChIP-Seq

Raw reads were quality-checked using FASTQC and adapters were trimmed using cutadapt. Reads were aligned to the hg19 genome using bowtie2^95^. Unmapped and low quality (MAPQ<10) reads were discarded. Duplicates were marked using Picard (https://broadinstitute.github.io/picard/) and removed using samtools^96^. From the replicate datasets, a consensus set of peaks were called for each TF at each stage using HOMER^97^ getDifferentialPeaksReplicates.pl using stage-matched input samples as background and -style factor. Briefly, first tag directories were created for each target and input replicate. Peaks were quantified for both target and input tag directories and DESeq2 was then invoked to identify peaks enriched in target ChIP samples over input using a fold enrichment threshold of 1.5 and fdr of 0.1. In the absence of replicates, peak calling was performed using macs2^98^ using –call-summits and a qvalue cutoff of 0.05. HOMER annotatePeaks.pl was then used to annotate these peaks.

Differential binding analysis was performed using the DiffBind package^99, 100^ using default parameters. Differentially bound ß-catenin and ATAC peaks were identified using a fold enrichment threshold of 1.5 and adjusted -pvalue < 0.05. A genomic site was defined as both ‘SOX17’ and TCF’ bound if: a significant binding event was called for both SOX17 and at least one of the TCF/LEF TFs using the relevant input sample as background, and no statistically significant differential binding between TFs was observed. A peak was called ‘SOX17 enriched’ if: the peak was called for only SOX17 and none of the TCFs, or if SOX17 binding displayed a greater log2 fold change over input relative to all the TCFs, and SOX17 was determined to have significantly increased binding over all TCFs quantified by DiffBind and DESeq2. Similarly, a genomic site was called ‘TCF-enriched’ if a peak was called for at least one of the TCFs but not SOX17, or at least one of the TCFs was determined to have significantly increased binding over SOX17. A fold enrichment threshold of 1.5 and FDR <0.1 to identify SOX vs. TCF enriched peaks, in order to also incorporate weakly bound TCF peaks. If a binding category contained less than 500 peaks, we didn’t use it for further analysis.

### ATAC-Seq

FASTQ files were quality-checked using FASTQC and Nextera adapters were trimmed using cutadapt. Trimmed paired-end reads were aligned to the hg19 genome using bowtie2 and the parameters -X 2000 -very-sensitive-local. Paired end bam files were filtered for mitochondrial reads, unmapped and low-quality reads. Duplicates were marked using Picard and removed using samtools. Peaks were called on replicates using Genrich (https://github.com/jsh58/Genrich) and the parameters j -r -e -v -q 0.1

### Nucleoatac analysis

As input for nucleoatac analysis, peak files of desired categories were extended upto 2000bp across peak summits. Nucleoatac^49^ was run using default parameters. For visualization and quantification of nucleosome occupancy, the occ.bedgraph files were converted to bigwigs using UCSC binary tools (https://hgdownload.cse.ucsc.edu/admin/exe/). Nucleosome occupancy scores were then computed using deeptools^101^ computeMatrix and visualized using plotProfile.

### Downstream data processing and visualization

To visualize ChIP-Seq and ATAC-seq data, filtered and sorted bam files were converted to bigwig files using deeptools bamCoverage with the parameters: -bS 20 –smoothLength 60 -e 200 –normalizeUsing RPGC using the effective genome size for GRCh37. Bigwig files were visualized using IGV^102^. Genomic algebra operations were performed using unix commands (awk, grep, sed) or using the bedtools suite^103^, particularly bedtools intersect to define overlapping genomic region of interest, or bedtools merge to define a union set of genomic regions. For all quantifications, merged bam files or bigwig files of both ChIP or ATAC replicates were used.

To identify patterns in ß-catenin time course ChIP-Seq datasets [Fig1, S1], an union of all ß-catenin peaks from all days was plotted using deeptools plotHeatmap and k-means clustering was performed using kclust 8. The most predominant 5 clusters were then extracted and retained for further analysis. Heatmaps, density plots or metaplots were generated using the deeptools package by invoking the computeMatrix (-reference-point center, -a 2500, -b 2500) and then plotHeatmap or plotProfile options. Volcano plots or MA plots of differential gene expression were generated using the EnhancedVolcano (https://github.com/kevinblighe/EnhancedVolcano) or ggpubr (https://github.com/kassambara/ggpubr) packages in R respectively. Heatmaps from RNA-Seq data was generated using the ‘pheatmap’ package.

Signal normalization (1/mapped tags/ sample such that each directory contains 10 million tags) and quantification was performed on merged ChIP/ATAC tag directories by HOMER. Boxplots and violin plots of ChIP/ATAC-Seq signal quantification were generated using the ggplot2 package and statistically significant differences in read density between conditions was determined by ANOVA or Wilcoxon rank sum test as appropriate in R. UpSET plots of the distribution of SOX17 or SOX2 and ß-catenin co-regulated enhancers were generated using *intervene*^104^. Data from ChIP-qPCR and reporter assays were visualized using GraphPad Prism. P-values were determined via nonparametric Mann-Whitney-U tests.

### DNA-binding Motif and GO enrichment Analysis

The MEME-Suite of tools^105, 106^ was used to perform *de-novo* motif analysis. For motif analysis, 100bp across peak summits were extracted for each category and converted to the fasta format using bedtools getfasta. *De-novo* motif analysis across peak sets was performed using DREME and default parameters, and motifs were identified using TOMTOM. Motif scanning at putative SOX17 or TCF-dependent enhancers for reporter assays was performed using FIMO with default parameters using the CIS-BP^107^ ‘Homo sapiens’ database as reference. GO term enrichment analysis was performed using GREAT^108^ and Gene Ontology^109, 110^.

### Analysis of public data

The following public datasets were reanalyzed as described above: GSM772971 (H3K4me1 ChIP-Seq in DE), GSM1112846, GSM1112844 (RNA-Seq of Day 3 ectoderm cells) and GSM1112835, GSM1112833 (RNA-Seq of Day 3 mesoderm cells)^43^.

---

**Figure S1 – Related to Figure 1.**
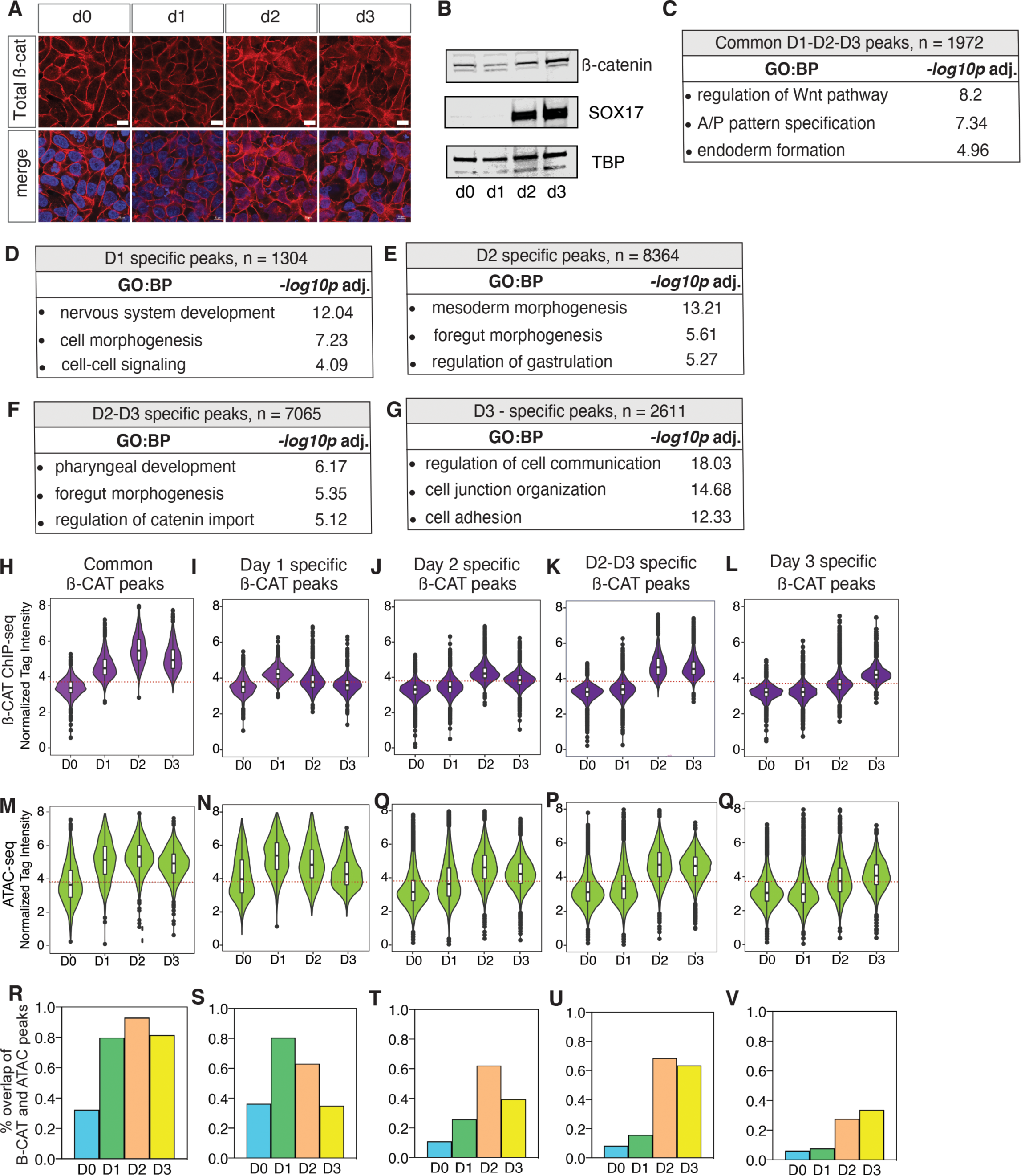
Characterization of ß-catenin chromatin binding during DE differentiation. **A.** Immunostaining showing total ß-catenin across days (d) 0 – 3 of differentiation (scale bar = 20 μm). **B.** Western blots of nuclear extracts showing total ß-catenin, SOX17 and TBP (loading control) proteins levels. **C - G.** GO enrichment analysis of five clusters of ß-catenin bound genomic regions: common peaks **(C)**, Day 1 specific peaks **(D)**, Day 2 specific peaks **(E)**, Days 2 -3 specific peaks **(F)** and Day 3 specific peaks **(G)**. For each category, the most enriched GO terms and the adjusted -log10 p-values (Fisher’s exact test, FDR 5%) are shown. **H – L.** Quantification of ß-catenin ChIP-Seq read densities of each category for each day. **M – Q.** Quantification of ATAC-Seq read densities. Dotted lines represent the approximate read density corresponding to the peak calling threshold. **H – K.** Statistical significance between read density of groups was determined by one-way ANOVA followed by multiple comparison via Tukey’s post-hoc honestly significant difference and results are available at Supplementary Table 3. *p <* 0.05 was considered significant in all cases. **R – V.** Bar graphs showing the percentage of ß-catenin peaks that overlap with ‘open’ ATAC-Seq peaks for each category at each day.

**Figure S2 – Related to Figure 1.**
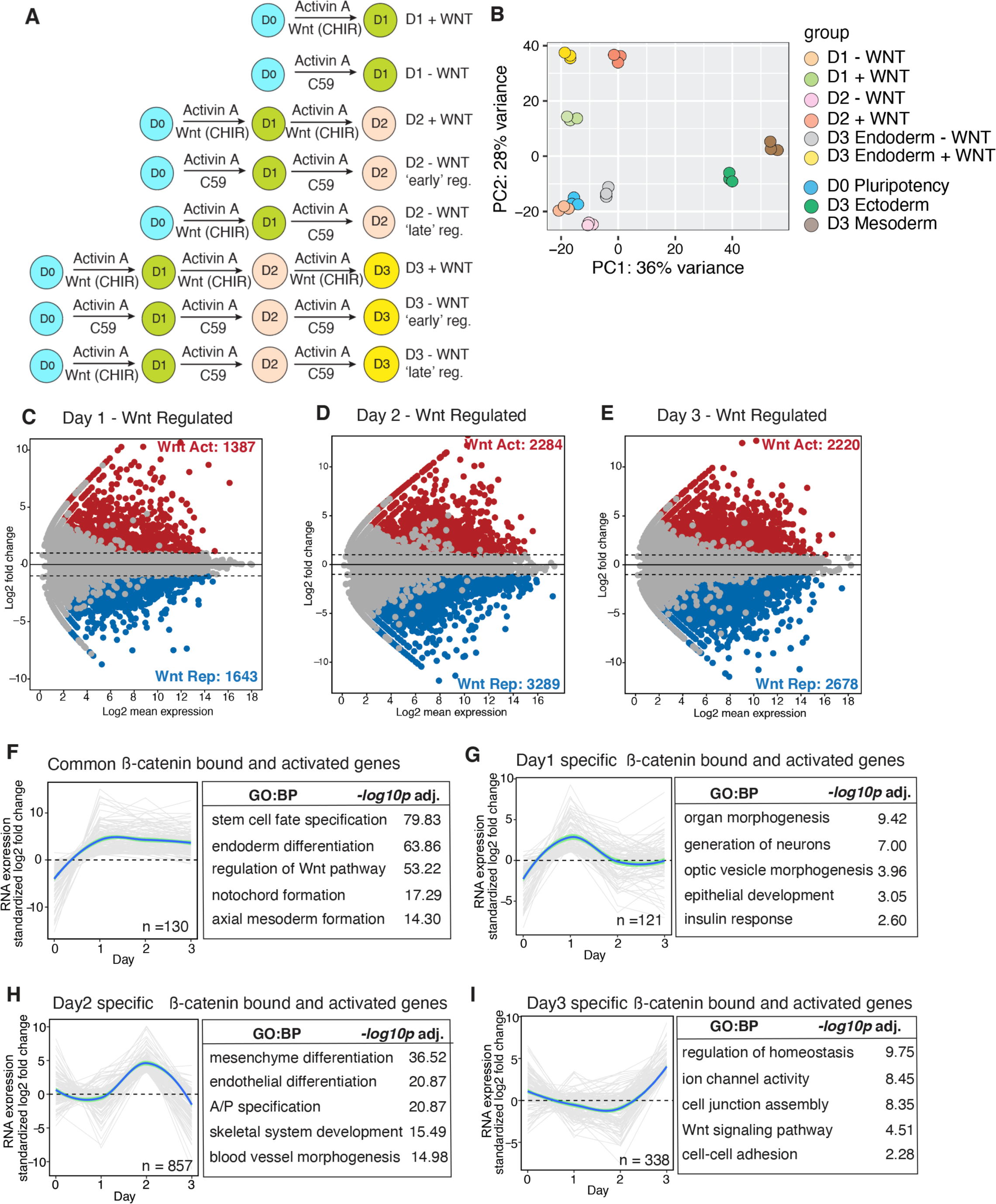
Dynamic Wnt-responsive genes during DE differentiation. **A.** Schematic showing different conditions and timings of CHIR and C59 treatment to identify Wnt regulated genes. **B.** principal component analysis (PCA) plot showing distribution of CHIR-treated (+WNT) and C59-treated (-Wnt) RNA-seq samples during endoderm differentiation, relative to day 3 mesoderm and ectoderm differentiation. **C - E** Differential expression analysis of +WNT versus -WNT samples (log2 fold change >1, p<0.05) to identify Wnt-responsive transcripts at days 1, 2 and 3 of differentiation. **F – I.** Relative expression levels of direct ß-catenin bound and Wnt activated genes plotted as a *loess* smoothed trendline (individual transcript data is shown in light grey) for the following categories: common genes bound and regulated by ß-catenin on all days of differentiation **(F)** Day 1 specific genes **(G)**, Day 2 specific genes **(H)** and Day 3 specific genes **(I)** and GO enrichment analysis of each category.

**Figure S3 – Related to Figure 2.**
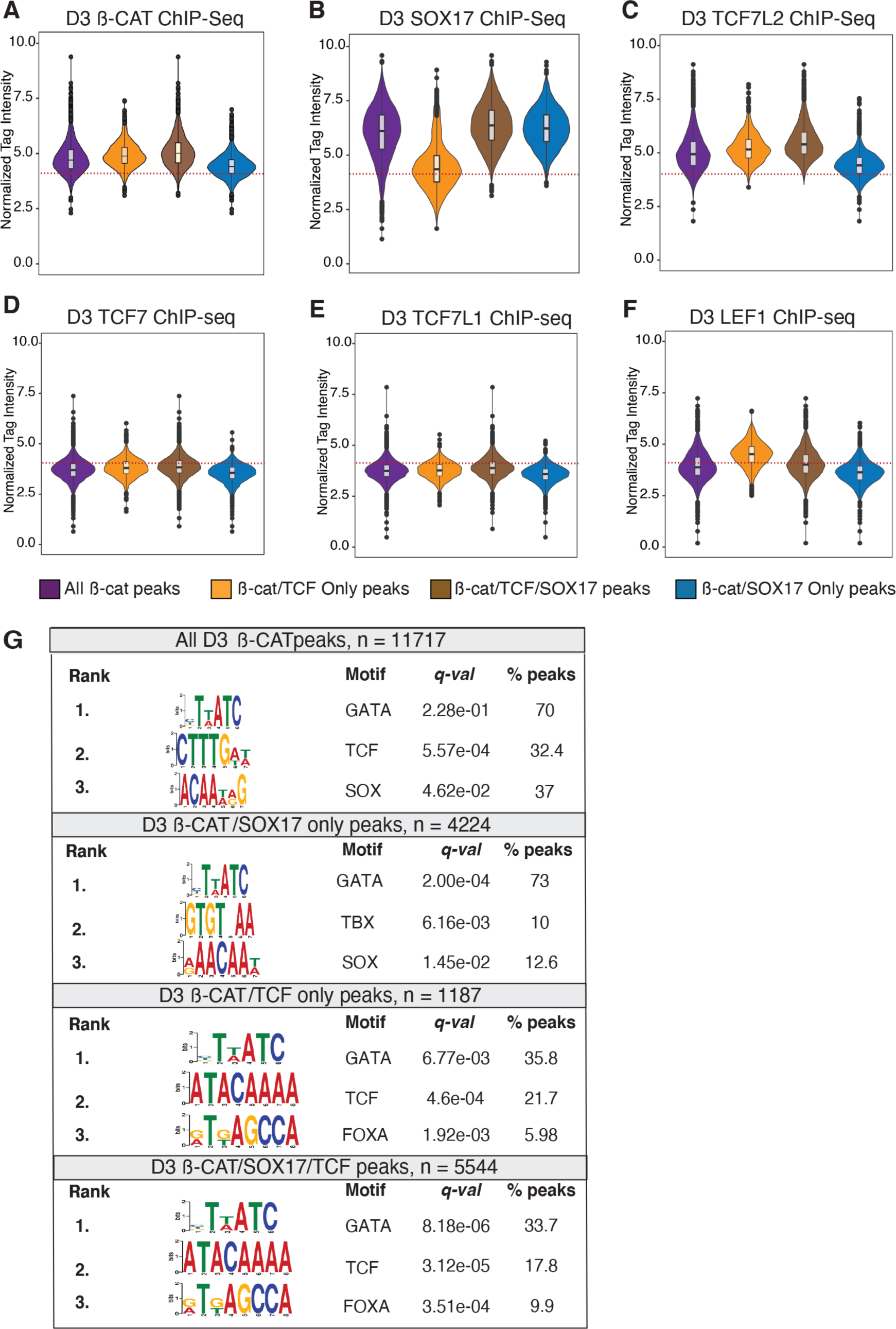
Differential co-occupancy of ß-catenin, SOX17 and TCFs in DE. **A – F.** Quantification of ChIP-seq tag density for ß-catenin **(A)**, SOX17 **(B)**, TCF7L2 **(C)**, TCF7 **(D)**, TCF7L1 **(E)** and LEF1 **(F)** at the following peak categories: All ß-catenin peaks, peaks bound only be ß-catenin and TCF. Peaks co-bound by ß-catenin, SOX17 and at least one TCF, peaks bound by ß-catenin and SOX17 but not TCFs. Dotted lines represent the approximate read density corresponding to the peak calling threshold. **G.** *De-novo* DNA-binding motif analyses of the above-described peak categories.

**Figure S4 – Related to Figure 3.**
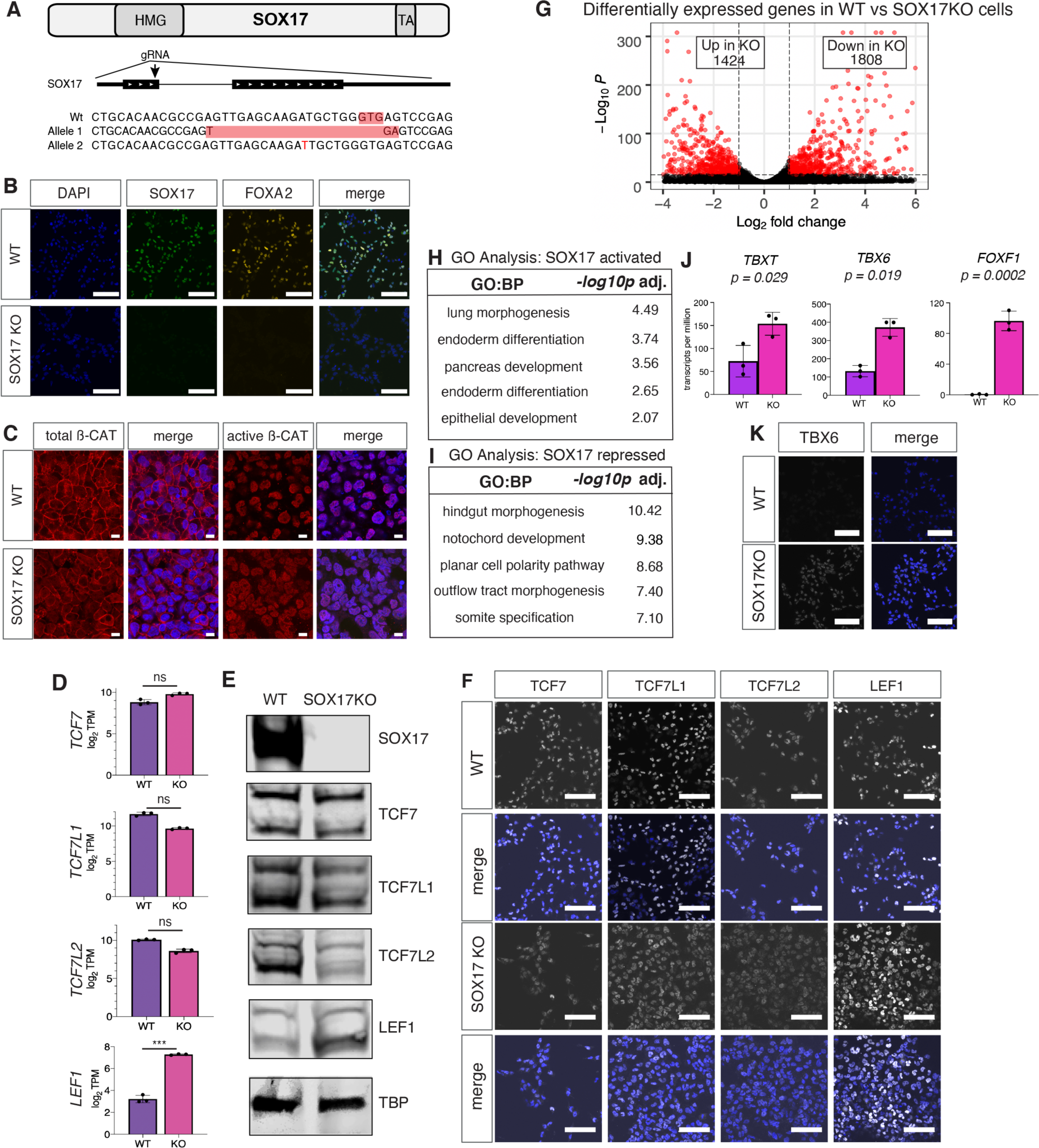
Characterization of SOX17 KO cells and identification of SOX17 regulated transcripts. **A.** Schematic showing the CRISPR-Cas9 targeting strategy to generate the homozygous mutant SOX17 knockout (KO) line. **B.** Immunostaining showing expression of endoderm markers SOX17 and FOXA2 in WT and SOX17KO cells (scale bar = 100 μm). **C.** Immunostaining of total and active ß-catenin protein levels in WT and SOX17KO cells (scale bar = 50 μm). **D.** RNA-seq expression levels (log2 transformed TPM) of *TCF7, TCFL1, TCF7L2* and *LEF1* in WT and SOX17KO cells. ***p= 0.0002, non-parametric Mann-Whitney U tests. Not significant (ns). **E.** Western blots and **F.** immunostaining showing protein expression levels of TCF7, TCF7L1, TCFL2 and LEF1 in WT and SOX17KO cells (scalebar = 100 μm). **G.** Volcano plot showing differentially ssed genes between WT and SOX17KO (Log2 foldchange >1, p< 0.05). **H– I.** Most enriched GO terms associated with SOX17 activated and SOX17 repressed genes and the adjusted -log10 transformed p-values (Fisher’s exact test, FDR 5%). **J.** RNA-seq expression levels (log2 transformed TPM) of exemplar mesoderm markers in WT and SOX17KO cells. P-values were calculated via the non-parametric Mann Whitney U test. **K.** Immunostaining of mesoderm marker TBX6 in WT and SOX17KO cells (scale bar = 100 μm).

**Figure S5 – Related to Figure 3.**
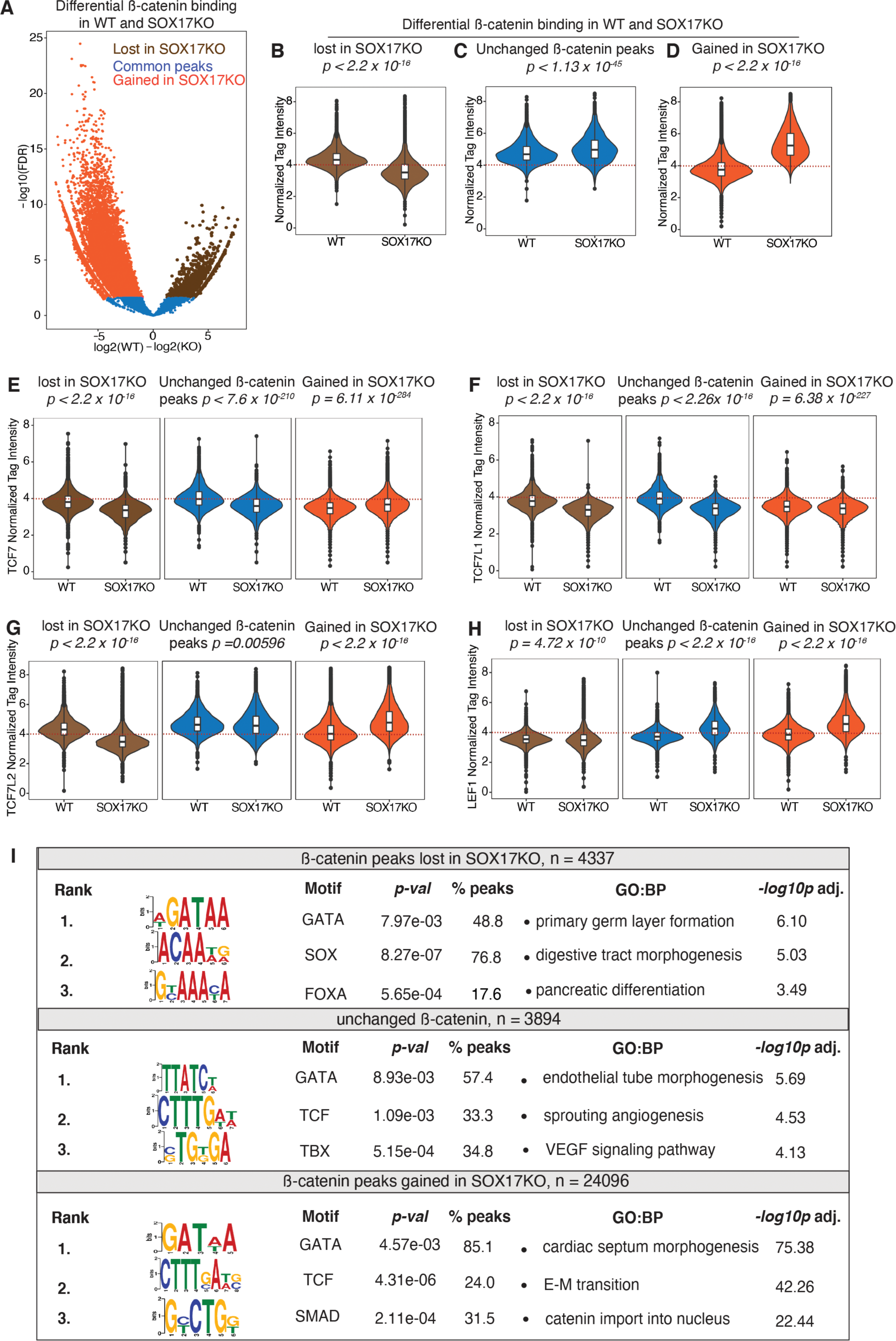
Differential ß-catenin ChIP-seq peaks in WT and SOX17 KO cells. **A.** Valcono plot showing distribution of differential ß-catenin binding events in WT and SOX17KO (fold change >1.5, FDR p< 0.05). Brown dots represent ß-catenin peaks lost in SOX17KO, blue dots are ß-catenin peaks gained in SOX17KO, and orange dots are ß-catenin peaks that did not change. **B - D.** Quantification of ß-catenin ChIP-seq density of each of the above-described categories in WT and SOX17KO cells plotted as boxplots. P values were calculated via the Wilcoxon rank sum test. **E – H.** Violin plots quantifying TCF7, TCF7L1, TCF7L2 and LEF1 ChIP-seq read densities at loci bound ß-catenin for each of the above categories in WT and SOX17KO cells. P values calculated by Wilcoxon rank sum test comparing WT and SOX17KO cells. **I.** *De-novo* DNA-binding motif analysis and GO term enrichment of genes associated with ß-catenin peaks in each category.

**Figure S6 – Related to Figure 3.**
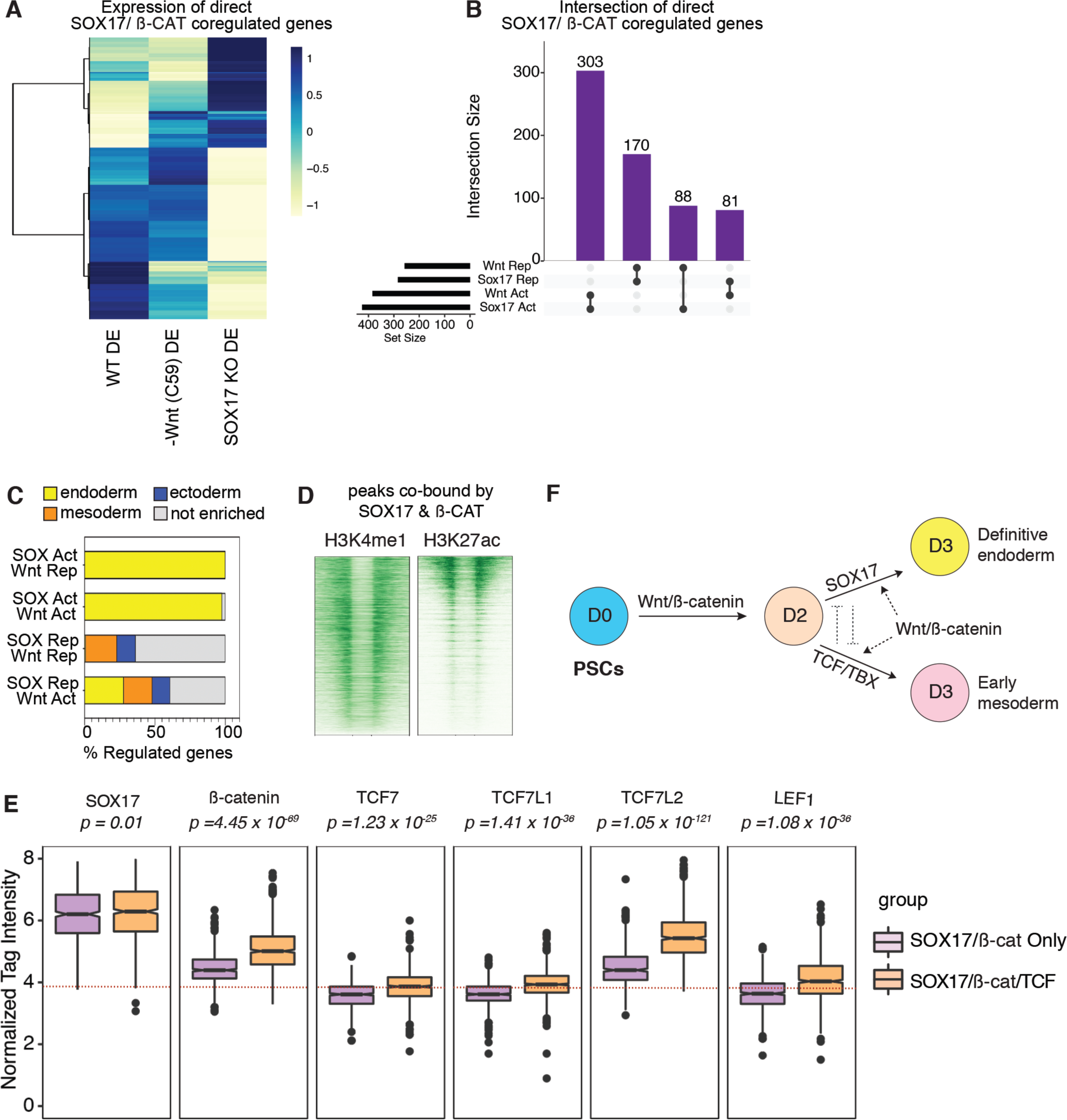
Analysis of genes co-regulated by SOX17 and WNT/ ß-catenin. **A.** Heatmap showing unsupervised clustering of RNA-seq expression for all genes cobound and coregulated by SOX17 and ß-catenin in wild type, C59 treated (-Wnt) or SOX17KO Day 3 DE cells. **B.** UpSET plot showing distribution of SOX17/ß-catenin coregulated genes that are also associated with co-bound enhancers, indicating whether a gene is activated (act) or repressed (rep) by SOX17 or WNT. **C.** Stacked bar graphs showing the percentage of genes associated with SOX17/ ß-catenin coregulated peaks that have enriched expressed in the entoderm (yellow), mesoderm (orange) or ectoderm (blue) peaks. Day 3 endoderm, mesoderm or ectoderm enriched genes were defined by re-analysis of the following datasets: GSM1112846, GSM1112844 (RNA-Seq of Day 3 ectoderm cells) and GSM1112835, GSM1112833 (RNA-Seq of Day 3 mesoderm cells) **D.** H3K4me1 (data analyzed from GSM772971) and H3K27ac ChIP-Seq density plots of 1670 loci cobound and coregulated by SOX17/ ß-catenin from Fig. 3G. **E**. Quantification of ChIP-seq read density for SOX17, ß-catenin, TCF7, TCFL1, TCF7L2 and LEF1 binding to loci that are occupied only by SOX17/ ß-catenin but not any TCF (pink) and loci occupied by co-occupied by SOX17, ß-catenin and at least one TCF. P values were calculated via the wilcoxon rank sum test. Dotted lines represent the approximate read density corresponding to the peak calling threshold. **F.** Schematic summarizing SOX17 and WNT/ß-catenin interactions during endoderm differentiation.

**Figure S7 – Related to Figure 4.**
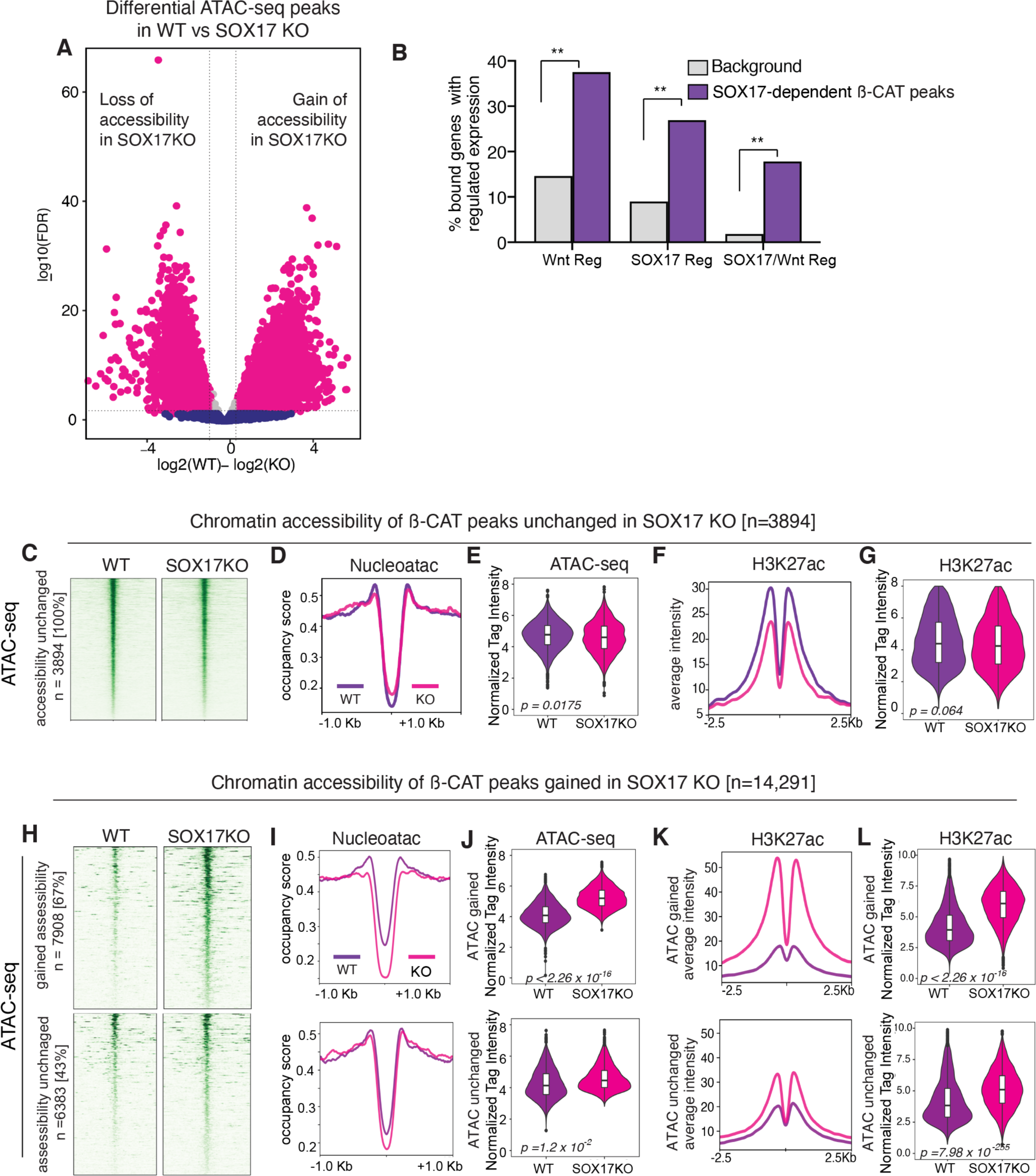
ATAC-seq and epigenetic analysis of ß-catenin bound loci. **A.** Volcano plot showing differential SOX17-dependent chromatin accessibility in WT and SOX17KO cells (fold change > 1.5 FDR p< 0.05) **B.** Bar graph showing the proportion of genes associated with SOX17-dependent ß-catenin chromatin binding and are not bound by TCFs, that have Wnt-regulated and/or SOX17-regulated expression. Background is all genes in the genome. ** *p* = 9.8 × 10^-208^ for Wnt regulated genes, *p* = 6.26 × 10^-187^ for SOX17 regulated genes, *p* < 2.2 × 10^-16^ for SOX17/Wnt coregulated genes, Fisher’s exact test. **C.** Density plot of ATAC-seq signal for ß-catenin peaks unchanged in SOX17KO. **D.** ATAC-seq metaplot showing average nucleosome occupancy signal at unchanged ß-catenin peaks and **E.** quantification of ATAC-Seq read densities. **F.** Metaplots showing averages H3K27ac ChIP-seq signal at loci with unchanged ß-catenin peaks and **G.** quantification of H3K27ac signal intensity as boxplots in WT and SOX17KO cells. P values calculated by the Wilcoxon rank sum test. **H.** Density plot of ATAC-seq signal for loci that gain ß-catenin peaks in SOX17KO cells. Heatmaps showing ATAC signal intensity of two classes of new ß-catenin peaks; those that gain accessibility in SOX17KO cells and those where accessibility is unchanged in both WT and SOX17KO. **I.** Metaplot of nucleoatac signal showing average nucleosome occupancy and **J.** quantification of ATAC-Seq signal at both classes of *de-novo* ß-catenin enhancers. **K.** Metaplot and **L.** quantification of H3K27ac ChIP-seq read intensity. P values were calculated via the wilcoxon rank sum test.

**Figure S8 – Related to Figure 5.**
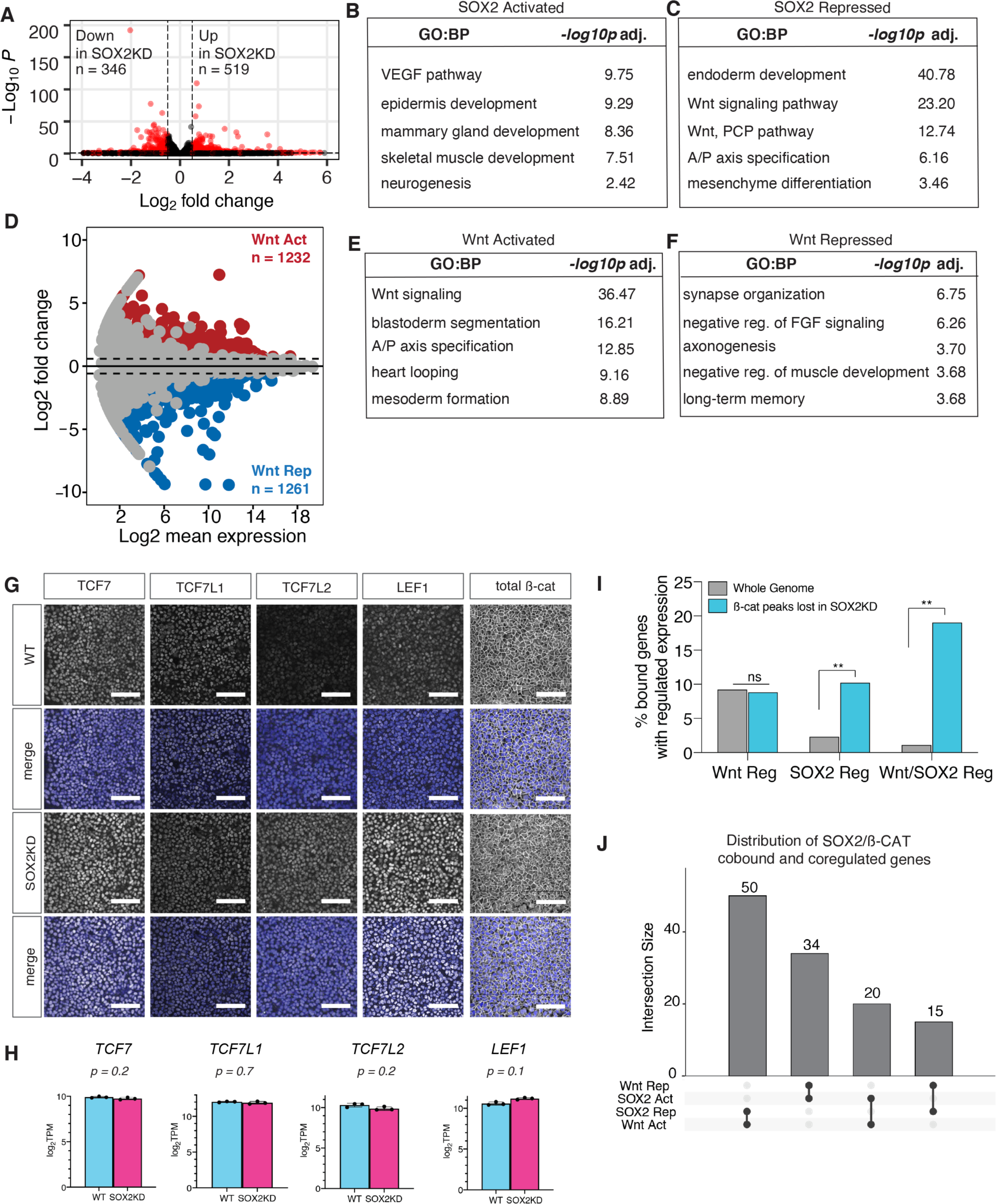
Identification of SOX2-regualted and WNT-regulated genes in NMPs. **A.** Identification of SOX2-regulated transcripts. Volcano plot showing differential gene expression (log2 fold change > 1, FDR p<0.05) upon CRISPR—mediated SOX2 knockdown (KD). **B – C.** GO term enrichment analysis of SOX2 activated and repressed genes. **D.** Identification of WNT-regulated transcripts in NMPs. MA-plot showing differential gene expression comparing Day3 NMPs differentiated with CHIR or C59 to inhibit WNT. **E – F.** Enriched GO terms associated with Wnt activated (Act) and Wnt repressed (Rep) genes, -log10 transformed p-values (Fisher’s exact test, FDR 5%). **G.** Immunostaining showing TCF7, TCF7L1, TCF7L2, LEF1 and total ß-catenin protein levels in WT and SOX2KD cells. (scalebar = 50 μm). **H.** RNA-seq expression levels (log2 TPM) of *TCF7, TCF7L1, TCF7L2* and *LEF1* in WT and SOX2KD cells. P-values were determined by two-tailed student’s T-test. **I.** Bar graph showing the proportion of genes associated ß-catenin peaks lost in SOX2KD that also have Wnt-regulated, SOX2-regulated and SOX2/Wnt-coregulated expression. Fisher’s exact test, *p* = 0.473 (ns) for Wnt regulated genes, *p* = 8.76 × 10^-29^ for SOX2 regulated genes, *p* = 7.10 × 10^-29^ for SOX2/Wnt coregulated genes. **J.** UpSET plot showing distribution of coregulated ß-catenin and SOX2 enhancers.

**Figure S9 – Related to Figure 5.**
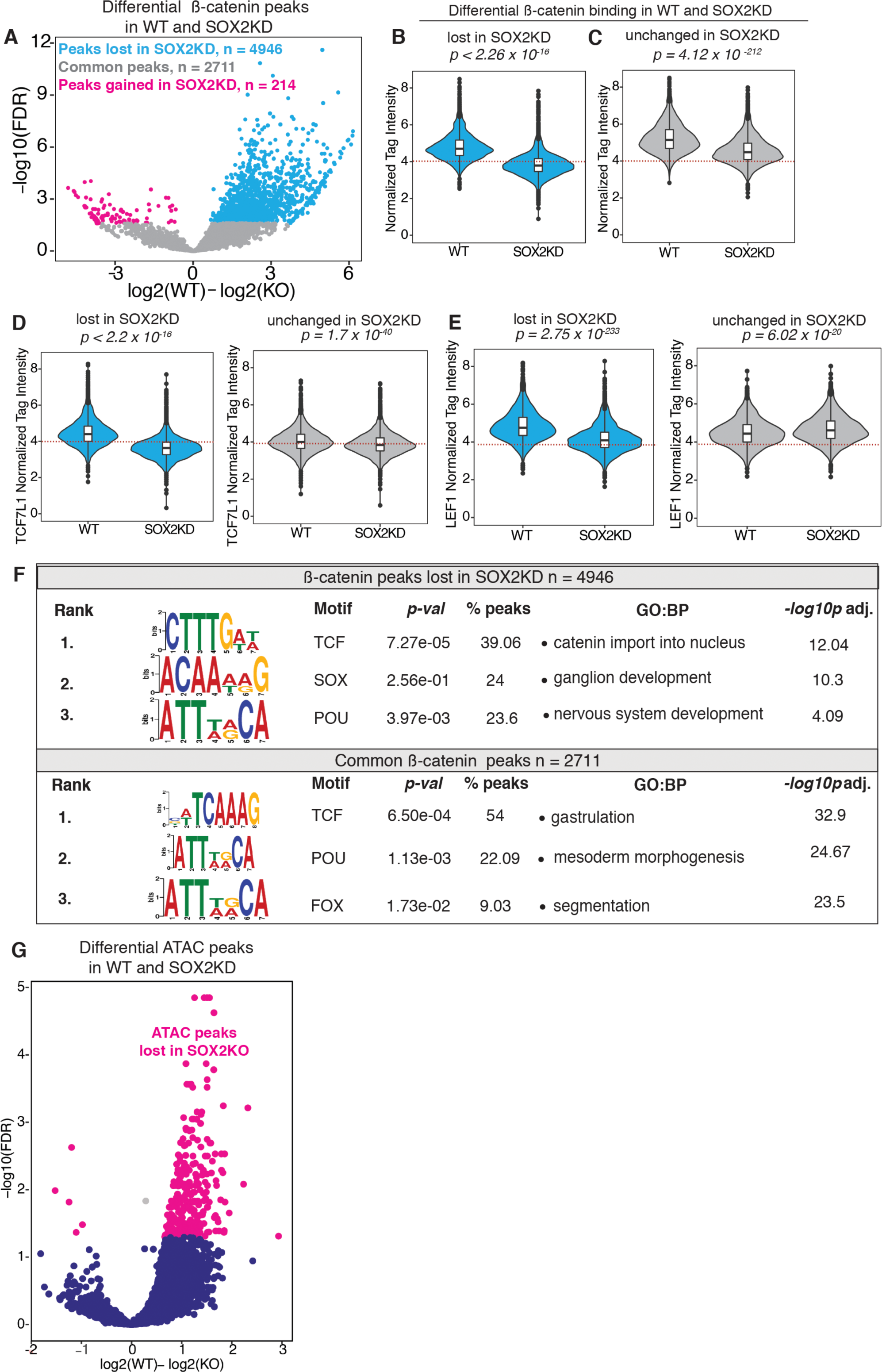
Analysis of SOX2-dependent ß-catenin chromatin binding. **A.** Volcano plot showing distribution of differentially bound ß-catenin ChIP-seq peaks in WT versus SOX2KD cells (fold change >1.5, p<0.05). **B – C.** Quantification of ß-catenin ChIP-seq read densities in WT and SOX2KD cells at loci that either lose ß-catenin (B) or where B-catenin binding is unchanged in SOX2KD. P-values were calculated via the Wilcoxon rank sum test. **D – E.** Quantification of TCF7L1 and LEF1 ChIP-seq read densities at both categories of peaks. P-values were calculated via the Wilcoxon rank sum test. Dotted lines represent the approximate read density corresponding to the peak calling threshold. **F.** Table showing *de-novo* motif DNA-binding analysis and GO term enrichment analysis of ß-catenin peaks that are lost or unchanged in SOX2KD NMPs. **G.** Volcano plot showing differentially accessible peaks as determined by ATAC-Seq in WT and SOX2KD cells (fold change >1.5, p<0.05).

**Figure S10 – Related to Figure 6.**
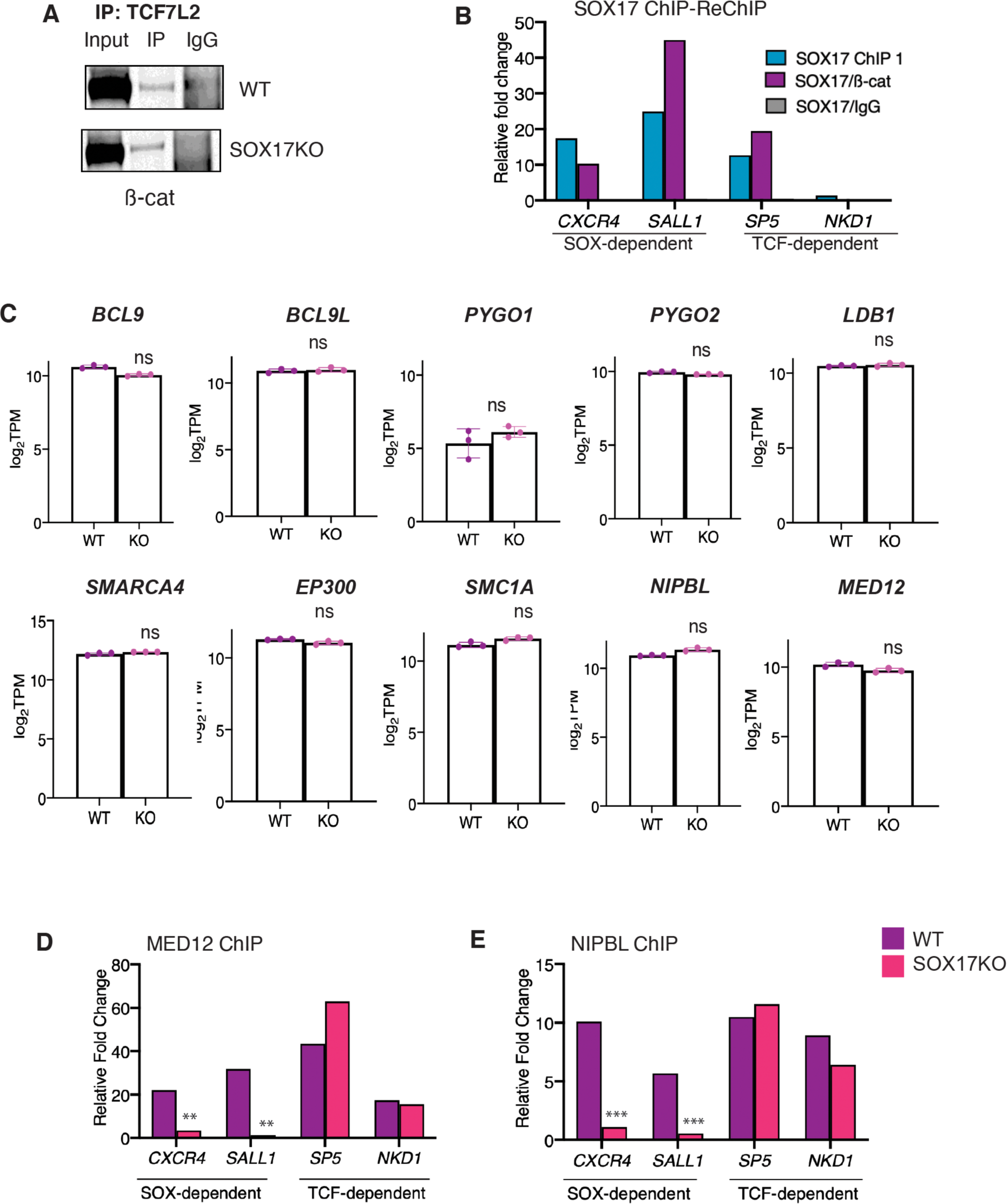
Analysis of enhancer complex components. **A.** Western blot confirming SOX17KO does not disrupt TCF7L2 and ß-catenin binding in co-immunoprecipitation. **B.** SOX17 ChIP-reChIP ys. Representative qPCR showing relative fold DNA recovery following either SOX17 ChIP-qPCR or ChIP-reChIP with ß-catenin at SOX-dependent or TCF-independent enhancers. **C.** RNA-seq expression levels (log_2_TPM) of Wnt-enhanceosome components or epigenetic interactors of ß-catenin in WT and SOX17KO cells. ns =not significant in two-tailed student’s T-test. **D – E.** ChIP-qPCR of MED12 and NIPBL showing relative DNA recovery at SOX-dependent or TCF-dependent enhancers. * = *p<0.05,* **=*p<0.01,* ***=p<0.001 based on two-tailed student’s T-test.

